# Large neutral amino acid levels tune perinatal neuronal excitability and survival

**DOI:** 10.1101/2022.07.12.499841

**Authors:** Lisa S. Knaus, Bernadette Basilico, Daniel Malzl, Maria Gerykova Bujalkova, Mateja Smogavec, Lena A. Schwarz, Sarah Gorkiewicz, Nicole Amberg, Florian Pauler, Thomas Rülicke, Jörg Menche, Simon Hippenmeyer, Gaia Novarino

## Abstract

Surprisingly little is known about the critical metabolic changes that neural cells have to undergo during development and how even mild, temporary shifts in this program can influence brain circuitries and behavior. Inspired by the discovery that mutations in *SLC7A5*, a transporter of metabolically-relevant large neutral amino acids, lead to a form of autism spectrum disorder, we employed metabolomic profiling to study the metabolic states of the cerebral cortex across different stages of life. We found that the cerebral cortex undergoes significant metabolic remodeling throughout development, with certain groups of metabolites showing stage-specific changes. But what are the consequences of interfering with this metabolic program? By manipulating Slc7a5 expression in neural cells, we found that the metabolism of large neutral amino acids and lipids in the cerebral cortex are highly interconnected. Deletion of *Slc7a5* in neurons perturbs specifically the postnatal metabolic state leading to a shift in lipid metabolism and a stage- and cell-type-specific alteration in neuronal activity patterns, resulting in a long-term cortical circuit dysfunction.

## INTRODUCTION

Human cortical development entails the timely coordination of a number of critical steps that are devised to generate a precise range of correctly positioned, connected and functional neuronal cells. These steps are guided and regulated by a network of genes whose mutations can underlie neurodevelopmental and neuropsychiatric disorders (Parenti et al., 2020). Challenging environmental conditions may also account for pathological variations of neurodevelopment but the identification of such factors is complicated and less understood (Galler et al., 2021; Rock and Patisaul, 2018; Ross et al., 2015; Stankovic and Colak, 2022). Genetic conditions, however, offer tractable entry points to isolate some of the extrinsic factors influencing the assembly of the brain.

We recently identified mutations in the gene *SLC7A5,* encoding a large neutral amino acid transporter (LAT1), as a rare cause of autism spectrum disorders (Tărlungeanu et al., 2016). Most of the large neutral amino acids are essential; thus, their presence in the human body is dependent on dietary intake. Currently, however, it remains largely unknown if and how the level of these amino acids changes over time in the brain and how fluctuations in their amount may influence the course of neurodevelopment. In fact, despite its importance, little is known in general about the metabolic program unfolding during brain development and the specific nutrient dependencies that this entails. For example, there is essentially no data on how brain metabolism changes immediately after birth. This critical period is especially interesting since the brain goes through a series of maturation processes while the organism has to adapt to new feeding and environmental conditions. Thus, understanding how specific nutrients can influence brain maturation may be key to prevent or correct aspects of certain neurodevelopmental conditions.

Here, we profiled the metabolome of the cerebral cortex at various developmental stages, defining significant longitudinal changes. Based on those results, we identified a perinatal time window when the forebrain exhibits an increased dependency on large neutral amino acids. Thus, we studied the effect of perturbing the perinatal metabolic state by limiting the amount of these essential amino acids in neural cells. In doing so, we identified a pivotal and unexpected function of large neutral amino acids during an exact temporal window crucial for cortical network refinement. Precisely, we found that altering the levels of large neutral amino acids in cortical neurons changes their lipid metabolism along with excitability and survival probability in a cell-autonomous manner, specifically early after birth. Our results offer a model of how mammalian neurons coordinate the expression of a nutrient-associated gene with the regulation of neuronal activity to ensure proper brain development. Changes in this program, during a very limited but critical time window, result in permanent cortical circuit defects.

## RESULTS

### Metabolome profiling reveals distinctive metabolic states of the cerebral cortex across development

Very little is known about the precise metabolic program adopted by the maturing brain. For example, previous studies have shown that neural progenitor cells (NPCs) mainly rely on anaerobic glycolysis (Figure 1A - left) (Bond et al., 2015; Candelario et al., 2013). On the other side, mature neurons meet their ATP demand by means of oxidative phosphorylation (Figure 1A -right), but, due to their high energy requirement and their inability to store glycogen (Duran et al., 2019; Vilchez et al., 2007), they rely on glial cells to provide metabolic support in the form of lactate (Mason, 2017; Philips and Rothstein, 2017). However, it is unknown how maturing neurons meet their energy demand at intermediate developmental stages when they lack full glial support (Figure 1A – middle).

**Figure 1.**
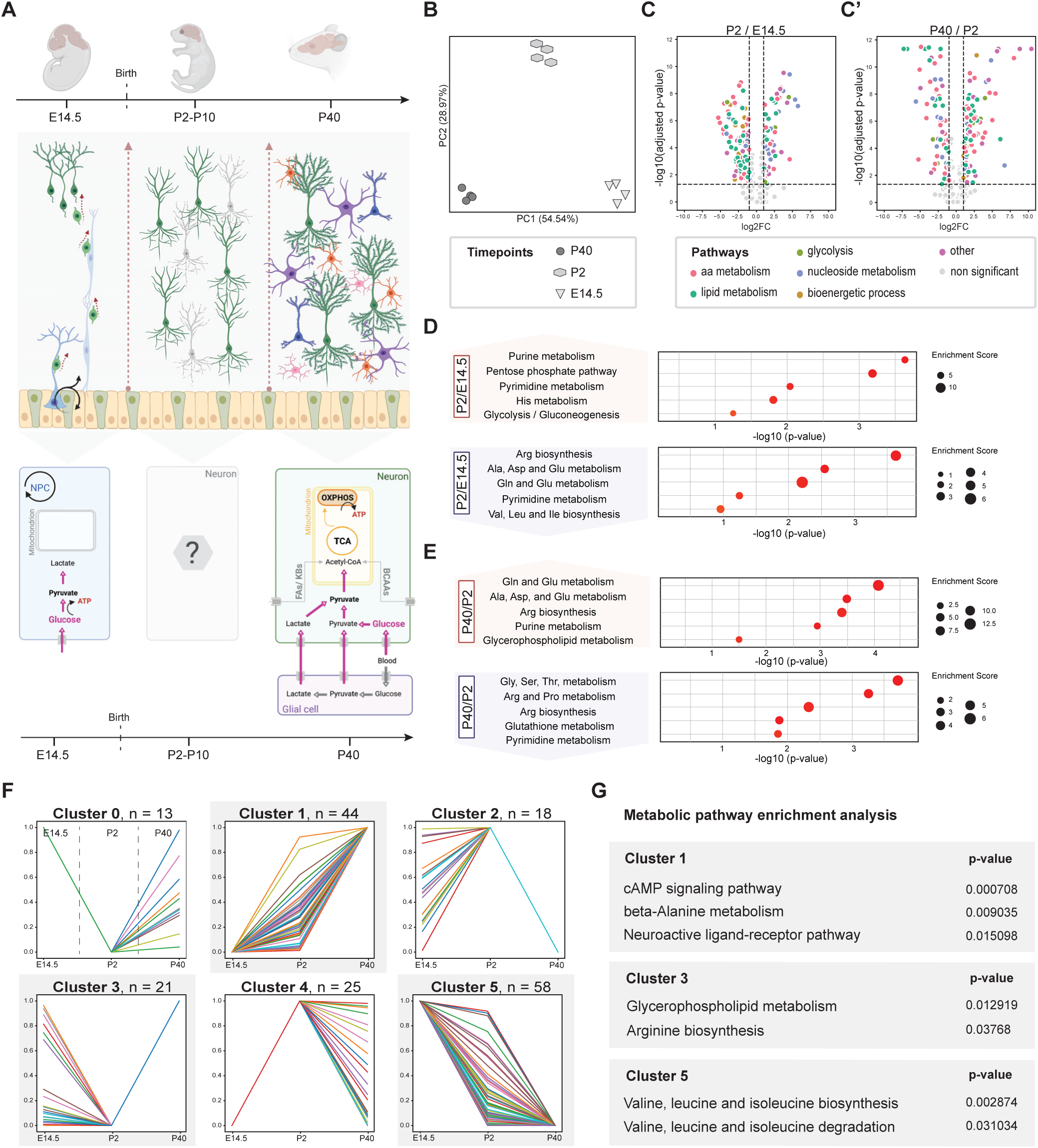
Metabolome profiling of the forebrain across time highlights developmental stage-specific metabolic states. (**A**) Schematic describing the ATP production strategies employed by neural cells in the cerebral cortex at different developmental stages. **Top left**: At E14.5, a proliferative neural progenitor pool gives rise to intermediate progenitors and a migratory population of immature excitatory neurons, which generate the different layers of the cortex. **Bottom left**: For ATP production, neural progenitors rely mainly on anaerobic glycolysis, a series of cytoplasmic biochemical processes converting glucose into lactate. **Top middle**: Perinatally (P2-P10), the immature cortical network undergoes major refinement. At this developmental stage, the majority of cell-types in the cortex are maturing excitatory and inhibitory neurons. Glial cells are detectable but immature. **Bottom middle**: The metabolic landscape of maturing neurons is largely unknown. **Top right**: Representation of a mature cortical network, including pyramidal (purple) and inhibitory neurons (blue) and glial cells (astrocytes (orange), oligodendrocytes (green) and microglia (pink)). **Bottom right**: Mature neurons utilize glucose as the main substrate for ATP production via aerobic glycolysis. Since the endogenously produced pyruvate levels are insufficient to meet mature neurons’ high energy demand, neurons are highly dependent on metabolic support, in the form of pyruvate or lactate, provided by glial cells. Pyruvate is channeled into the mitochondrial ATP production pathways, including the TCA cycle and oxidative phosphorylation (OXPHOS). (**B**) Principal component analysis of the metabolome of E14.5, P2 and P40 cortices of wild-type mice. (**C-C’**) Volcano plot showing differentially abundant metabolites in wild-type animals across developmental stages. Metabolites were manually annotated to five main metabolic pathways. (**D-E**) Metabolic pathway enrichment analysis of all significantly up-(**top**) or down-regulated (**bottom**) metabolites at (**D**) P2 compared to E14.5 and (**E**) P40 compared to P2. (**F**) Clustering of KEGG-annotated metabolites based on their trajectory over time; (x-axis: age; y-axis: scaled abundance). (**G**) Metabolic pathway enrichment analysis revealed an overrepresentation of metabolites related to ‘cAMP signaling pathway’ (*p<0.00075*; *adj-p<*0.076), ‘beta-Alanine metabolism’ (*p<0.0095*; adj-p<0.12) and ‘Neuroactive ligand-receptor pathway’ (*p<0.02*; *adj-p<*0.27) in cluster 1. Cluster 3 is enriched for the metabolic terms ‘Valine, leucine and isoleucine biosynthesis’ (*p<0.003*; *adj-p*<0.31) and ‘-degradation’ (*p<0.0032*; *adj-p=1*). Cluster 4 is enriched for pathways including ‘Glycerophospholipid metabolism’ (*p<0.013*; *adj-p<*0.73) and ‘Arginine biosynthesis’ (*p<0.038*; *adj-p<*1; Supplementary data 2-7).

To gain an understanding of the metabolic states and transitions occurring during brain maturation, we performed a metabolomic analysis of the mouse cerebral cortex obtained at three different time points: embryonic day 14.5 (E14.5), postnatal day two (P2), and postnatal day 40 (P40). These time points coincide with different feeding strategies and are enriched for neural cells in different states, that is, neural progenitors and immature (E14.5), maturing (P2), and mature (P40) neurons (Figure 1A). By employing two independent High Performance Liquid Chromatography (HPLC) detection strategies (i.e., Hydrophilic interaction chromatography (HILIC) and Reverse-phased chromatography (RP)), we quantified a total of 346 metabolites. Principal component analysis of the results clearly separated the samples based on sampling age (Figure 1B), indicating that the cerebral cortex at each of these three time points is in a distinct metabolic state. The metabolic reorganization occurring during the development and maturation of the cortex is very extensive, with 273 out of 346 metabolites showing significant changes over time (Figure 1C-C’). The variability across samples was relatively small, with most of the changes being comparable across animals (Figure S1A), indicating that these changes are tightly regulated. Specifically, compared to E14.5, at P2, the level of 137 metabolites is significantly decreased and only that of 60 significantly increased (Figure 1C). At P40, 202 metabolites show a significantly different level compared to P2, with the changes being approximately equally divided between increased and decreased levels (Figure 1C’). Enrichment analysis revealed an overrepresentation of purine- and pentose phosphate-related metabolites among the metabolites increasing at P2 (compared to E14.5) and a significant enrichment of amino acid-related metabolites among those decreasing at the same stage (Figure 1D). At P40, instead, we detected a substantial increase in glutamine and glutamate-related metabolites and a decrease in a different set of amino acids (Figure 1E), disclosing a stage-specific regulation of amino acid metabolism. To better understand the quality of those changes over time, we plotted the developmental trajectories of all the detected metabolites (n= 346; Figure S2, Suppl. data 1). Next, employing a Gaussian mixture model (GMM), we clustered all KEGG annotated metabolites (n=176) exhibiting similar developmental trajectories (Figure 1F) and assessed whether the different clusters were enriched for particular classes of metabolites (Figure 1G). This analysis revealed a predictable enrichment for metabolites belonging to neuroactive and cAMP signaling pathways in the cluster representing metabolites with ascending trajectories over time (Figure 1F-G - cluster 1), with the majority of the metabolites steeply increasing between P2 and P40. On the other hand, we found that cluster 5, the cluster of metabolites showing declining trajectories over time, is enriched for branched-chain amino acids (BCAAs) and BCAA-related metabolites (Figure 1F-G - cluster 5). This cluster particularly sparked our interest since the BCAAs are substrates of the SLC7A5 transporter. We reasoned that the decline over time of these and other few amino acids (i.e., Figure S2, metabolite (M) 016, M018, M032, M077, M094) could indicate either a decreasing brain intake or an increasing utilization by neural cells as they mature. Either way, this observation further suggests that neural cells have significantly distinctive amino acid demands at different developmental stages. Finally, the only other cluster enriched for specific classes of metabolites is cluster 3, which displays a drop of glycerophospholipid- and arginine-related metabolites specifically at P2 (Figure 1F-G - cluster 3). To further expand our analysis, we performed a literature-curated annotation of all the detected metabolites (including the ones without KEGG-annotation), assigning them to six main metabolic pathways (i.e., amino acid (AA), lipid and nucleoside metabolism, bioenergetic processes, glycolysis and other; Figure 1C-C’). Thus, we repeated the clustering and enrichment analysis employing all the metabolites and our annotations (Figure S1B-B’). This analysis confirmed an enrichment of lipid and amino acid-related metabolites in clusters that show, respectively, a specific drop and a steep decrease at P2.

Of note, several of the metabolites we detected and that dramatically change concentration over time, are regulated by neurodevelopmental disease-associated genes (Figure S1C). For example, loss of the gene *PRODH* in humans causes hyperprolinaemia, associated with a syndromic form of epilepsy, mental retardation and autism spectrum disorder (ASD) (Afenjar et al., 2007; Di Rosa et al., 2008). Further, genes coding for enzymes involved in hydroxyproline (*P4HA2*) and guanine (*GDA*) metabolism (De Rubeis et al., 2014; Li et al., 2014) as well as *AMPD1* (Xia et al., 2014; Zhang et al., 2015), a gene involved in the purine nucleotide cycle, are all considered ASD high-risk genes. Thus, our dataset may be useful to identify developmental time windows underlying the onset of neurological conditions associated with abnormal regulation of brain-relevant metabolites.

### Perturbation of large neutral amino acid uptake leads to perinatal disruption of lipid metabolism

With a better understanding of the metabolic landscape of the cerebral cortex and the notion that the level of the BCAAs and the BCAA-related metabolites does significantly change during brain maturation, we moved on investigating the consequences of a decreased BCAA supply during this critical period. Therefore, we specifically ablated *Slc7a5*, the main BCAA transporter, in neural cells as an entry point to study the function of these amino acids in brain development. Indeed, although Slc7a5 has been primarily described as a blood-brain barrier transporter (Tărlungeanu et al., 2016), we found that it is also expressed in neural cells, particularly perinatally (Figure 2A-B), coinciding with the time window displaying a drop in BCAA levels in the mouse cerebral cortex (Figure 1F-G - cluster 5). In addition, while patients with *SLC7A5* mutations present with severe microcephaly, in mice the deletion of *Slc7a5* from the blood-brain barrier (*Slc7a5*^fl/fl^;*Tie2-*Cre positive (+) mice) does not lead to a reduction in brain size (Tărlungeanu et al., 2016). This suggests a function of *SLC7A5*, and its substrates, in cell types other than the endothelial cells of the blood-brain barrier. Thus, we crossed floxed *Slc7a5* mice (*Slc7a5^fl^*) with *Emx1-*Cre animals. *Emx1-*Cre mice express the Cre recombinase in the radial glial cells of the dorsal telencephalon from E9.5 on (Gorski et al., 2002), thereby inducing *Slc7a5* deficiency in NPCs and their progeny, including the excitatory neurons of the neocortex and hippocampus as well as in the glial cells of the pallium (Figure 2A and Figure S1D). Deletion of *Slc7a5* in neural cells reduces *Slc7a5* expression levels in the cerebral cortex by 50% (Figure S1E), an expected partial reduction considering that *Slc7a5* expression at the blood-brain barrier and in interneurons is not affected in the *Slc7a5*^fl^;*Emx1-*Cre mouse line.

**Figure 2.**
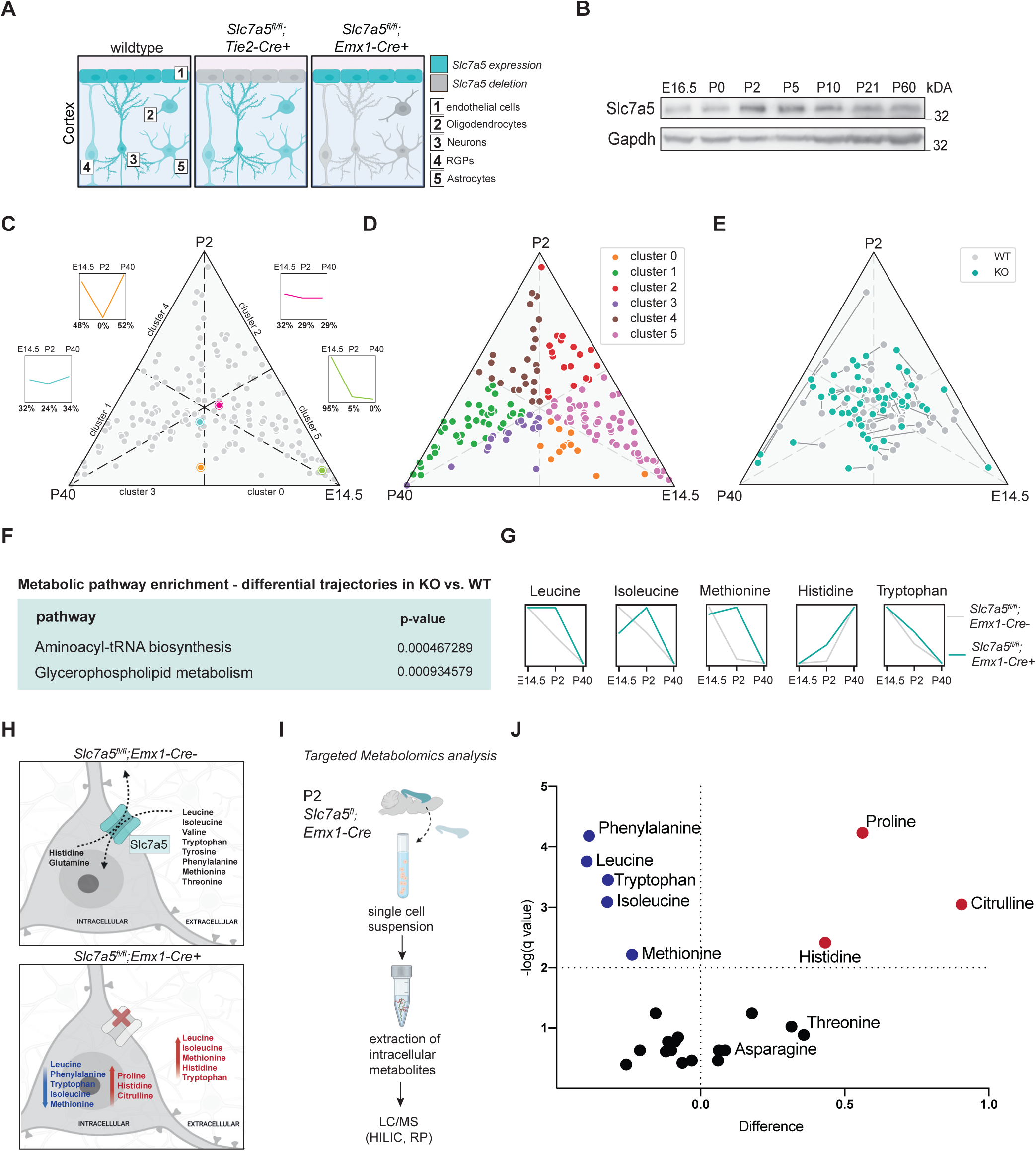
The neonatal metabolic state is dependent on *Slc7a5* expression. (**A**) Schematic representation of *Slc7a5* expression in different cell-types in the cerebral cortex of a wild-type (**left**), *Slc7a5^fl/fl^;Tie2-Cre+* (**middle**) or *Slc7a5^fl/fl^;Emx1-Cre+* (**right**) mouse. (**B**) Western blot analysis of *Slc7a5* expression in cortical samples obtained from *Slc7a5^fl/fl^;Tie2-Cre+* animals over the course of development. (**C-E**) Ternary plot classification of metabolites in the wild-type and mutant cortex. (**C**) The localization of each metabolite (dot) within the ternary plot is determined by 1) its previously determined cluster affiliation (Figure 4B) and 2) its exact unscaled trajectory, which is defined as the ratio between the three different time points (E14.5, P2 and P40). (**D**) Ternary plot of all KEGG-annotated metabolites in control animals. (**E**) Compared to wild-types (grey dots), the loss of *Slc7a5* leads to changes in the trajectory shape of a large fraction of metabolites (cyan dots). (**F**) Metabolic pathway enrichment analysis of all the KEGG-annotated metabolites whose trajectory over time was affected by *Slc7a5* deletion (Supplementary data 9). (**G**) The loss of *Slc7a5* leads to a stage-specific accumulation of the transporter’s substrates in cortical tissue (n=4 animals per genotype and time point; Pearson’s coefficient: Supplementary data 1; x-axis : age; y-axis: scaled abundance). (**H**) Slc7a5 facilitates the flux of BCAAs and other large neutral amino acids across the neuronal membrane (**top**). Loss of *Slc7a5* causes an extracellular accumulation and an intracellular depletion of BCAAs and some large neutral amino acids (**bottom**). (**I**) Experimental workflow employed for the quantification of targeted intracellular metabolites in cortical samples obtained from 2-days old *Slc7a5^fl/fl^;Emx1-Cre* mice. (**J**) Volcano plot of all the canonical amino acids measured in *Slc7a5* mutant and control cells. Amino acids showing significantly lower (blue) or higher (red) levels in *Slc7a5^-/-^* cells are indicated (n = 7 per genotype; male and female littermate pairs; FDR cut-off: 1.0%).

Next, we analyzed the metabolome of the cerebral cortex of *Slc7a5*^fl/fl^;*Emx1*-Cre+ mice and compared the profile of mutant and wild-type tissue over time. This comparison offers the opportunity to resolve i) the cause of the drop in large neutral amino acids (particularly BCAAs) observed in P2 wild-type cortical lysates (Figure S2, metabolites M016, M018, M032, M077, M094) and ii) the impact of deregulating these amino acids on the metabolism of the cerebral cortex. We found that the effect of *Slc7a5* deletion on the overall metabolite profile is relatively minimal (Figure S1F). Thus, to investigate whether *SLC7A5* mutations affect specific groups of metabolites, we performed an enrichment analysis of the KEGG-annotated metabolites showing divergent trajectories (r < 0.975) in mutants and controls (Figure 2C-E, Figure S3, Suppl. data 1). As expected, this analysis identified the pool of amino acids transported by Slc7a5, grouped into the metabolic term ‘Aminoacyl-tRNA synthesis’ (Figure 2F), as the top class of affected metabolites in mutant animals. We were surprised to find, however, that the deletion of *Slc7a5* affects the level of these amino acids in cortical tissue only at P2 (Figure 2G), implicating an important function of Slc7a5 specifically in neonatal mice. Furthermore, we noticed that the level of these amino acids is higher, not lower, in mutants compared to controls (Figure 2G). This observation suggests that in the absence of Slc7a5, the large neutral amino acids accumulate in the extracellular space and are not consumed by neural cells (Figure 2H). To test this possibility, we quantified amino acid levels in neural cells isolated from P2 control and mutant cortical tissue (Figure 2I), thereby measuring their intracellular amount. Indeed, we found that intracellularly the level of the primary Slc7a5 substrates is significantly reduced in mutant cells. In contrast, the level of histidine, the counter amino acid (Napolitano et al., 2015), is increased (Figure 2J). Thus, in the absence of Slc7a5, the large neutral amino acids are not transported and used inside neural cells, specifically at P2. These results also indicate that the drop in amino acid levels observed in P2 control samples (Figure 2G) reflects an increased utilization by neural cells rather than a reduced demand.

Interestingly, despite that the BCAAs are a source of Acetyl-CoA, an essential compound for the TCA cycle (Ye et al., 2020), we observed only a very mild reduction in the intracellular levels of some of the key players of energy storage and transfer (Figure S4A-F). In agreement, while a derailment of energy homeostasis has been associated with increased levels of oxidative stress (Aon et al., 2010; Murphy, 2009; Robb et al., 2018), in *Slc7a5* mutant cells, the ratio between reduced and oxidized Glutathione was not altered (Figure S4G). Surprisingly, however, we identified another group of metabolites that is strongly influenced by the loss of *Slc7a5* expression in neurons that are the metabolites related to ‘glycerophospholipid metabolism’ (Figure 2F), the same metabolites showing a sharp transition at P2 in control cortices (Figure 1G – cluster 3), suggesting a connection between Slc7a5-transported amino acids and lipids.

Next, we aimed to identify potential signaling cascades responding to this shift in metabolite profile. Remarkably, we found that the decrease in intracellular large neutral amino acids does not lead to the derailment of pathways associated with amino acid sensing and protein synthesis defects, such as the mTOR, the AMPK, or the unfolded protein response (UPR) signaling cascades (Figure S5)(Iurlaro and Muñoz-Pinedo, 2016; Nwadike et al., 2018; Riggs et al., 2005; Takahara et al., 2020; Wortel et al., 2017; Zhang et al., 2021). Thus, to uncover alternative consequences of the observed metabolite profile, we performed a comparative proteomic study of neonatal control and mutant cerebral cortices. Our analysis resulted in the detection of ∼ 8000 protein groups which were filtered based on fold change (comparing mutants to wild-type littermates) and False Discovery Rate (FDR) thresholds. Applying an FDR threshold of 1%, we identified 1202 deregulated proteins, comprising 954 up-regulated and 248 down-regulated proteins in the *Slc7a5^fl/fl^;Emx1-Cre+* cortex (Figure 3A). Thus, we employed Gene Ontology GO enrichment analysis to identify biological processes and cellular components that are significantly affected by *Slc7a5* deletion. This analysis returned proteins involved in lipid metabolism as being the most significant and numerous among the up-regulated proteins (Figure 3B - right). While, among the down-regulated GO-terms, we found an enrichment for neuron projection and membrane-associated proteins (Figure 3B - left). Those GO-Terms were reproducible at the mRNA expression level (Figure S4H).

**Figure 3.**
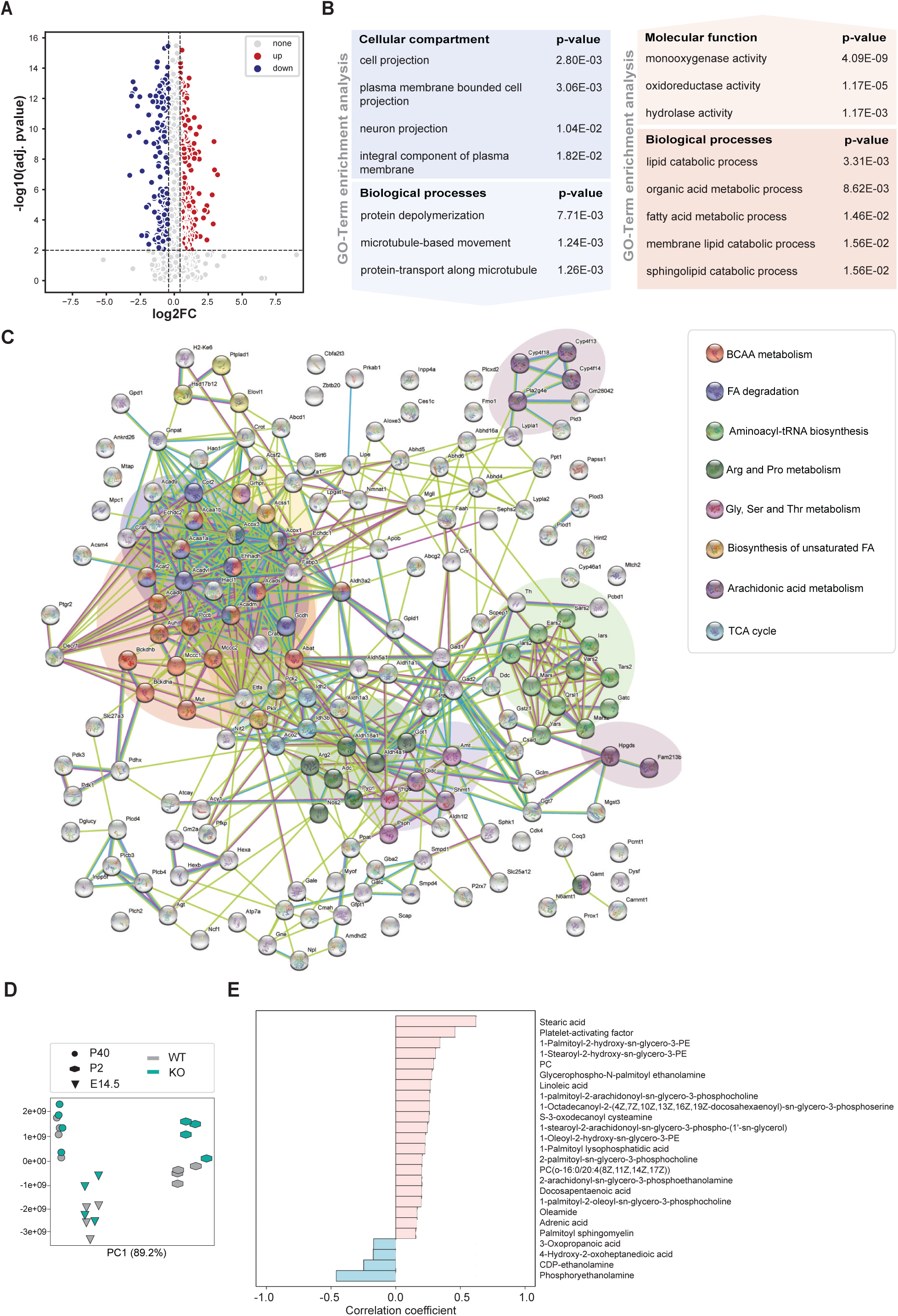
Branched-chain amino acid deprivation alters neuronal lipid metabolism. (**A**) Volcano plot of deregulated proteins at 1% FDR cut-off in the *Slc7a5^fl/fl^;Emx1-Cre+* cortex at P5. (**B**) GO-term enrichment analysis of the up-regulated and down-regulated proteins at 2% FDR of the *Slc7a5^fl/fl^;Emx1-Cre+* cortex (selected GO-terms: Supplementary data 10-11). (**C**) Protein-protein interaction network (String network) of all the proteins up-regulated (FDR<1%) in the *Slc7a5^fl/fl^;Emx1-Cre+* cortex. (**D**) PCA plot of all fatty acids (FA) and lipids detected in wild-type and mutant mice over time using an untargeted metabolomics approach (n=4 animals per genotype and time point). (**E**) Top 25 lipids/FAs correlating with the mutant genotype.

Next, we performed a protein-protein interaction analysis to spot potential connections between the large neutral amino acids and lipid-associated proteins. This analysis not only indicated an enrichment for BCAA- and lipid-related proteins among the proteins up-regulated in mutant animals, but also disclosed a direct interaction between these two classes of deregulated proteins (Figure 3C). Considering this strong enrichment for lipid-relevant terms in the proteomic dataset, we went back to the metabolomic dataset to check how the level of various fatty acids and lipids is affected by *Slc7a5* mutations. First, we found that the lipid profile of the cortex dramatically changes throughout development (Figure 3D), suggesting an active role of lipid composition in brain development. Second, we noticed that while at E14.5 and P40 mutants and control samples cluster together, P2 samples separate by genotype (Figure 3D), thereby underlining a timepoint-specific alteration in lipid composition due to the loss of *Slc7a5*. To follow this up, we assessed which lipids are either positively or negatively correlated with *Slc7a5* deletion over time (Figure 3E). Among these lipids showing a positive correlation with the mutant genotype, a few stood out for their role in neurons. These include, for example, the platelet-activating factor, an active phospholipid implicated in various neuronal functions (Clark et al., 1992; Hammond et al., 2015) whose level drops in control cerebral cortices only at P2 (Figure S2, metabolite M014) and the docosapentaenoic acid (Dyall, 2015). Altogether, these findings suggest an increased perinatal dependency of neural cells on large neutral amino acids. Decreased availability of these amino acids does not lead to a strong derailment of cellular metabolism but reveals a direct link between branched-chain amino acid and lipid metabolism in the brain.

### Lack of the amino acid transporter SLC7A5 leads to stage-specific neuronal cell loss

Can the perinatal metabolic shift displayed by *Slc7a5* mutant neural cells explain some clinical phenotypes reported in patients? *Slc7a5*^fl/fl^;*Emx1-*Cre+ mice are born at Mendelian ratio, are viable and at birth do not display obvious growth defects compared to their wild-type littermates (i.e., *Slc7a5*^fl/fl^;*Emx1*-Cre- or *Slc7a5*^fl/+^;*Emx1-* Cre+ mice) (Figure S6A). In agreement with the amino acid profile that changes only postnatally, *Slc7a5*^fl/fl^;*Emx1*-Cre+ mutants are born with normal brain size (Figure S6B-C’), indicating that *Slc7a5* is dispensable during the proliferative phase of the NPCs of the forebrain. Thus, *SLC7A5* mutations do not lead to microcephaly by hindering the generation of an appropriate number of neurons. However, by P40, the brain of *Slc7a5*^fl/fl^;*Emx1-*Cre+ mice is considerably smaller than that of their control littermates (Figure 4A-A’). In agreement, histological analysis of mutant and wild-type brains revealed a severe reduction in the thickness of the cerebral cortex in P40 *Slc7a5*^fl/ fl^;*Emx1-*Cre+ animals (Figure 4B-B’), with layers II-III being the main drivers of this difference (Figure S6D-F’). To identify a potentially critical time window for this phenotype to emerge, we monitored the brain size of mutant and wild-type littermates throughout pre- and postnatal development. Interestingly, the difference in brain size between control and *Slc7a5* mutant animals appears during the first postnatal week (Figure 4C) and remains stable from P10 on, coinciding with the increased *Slc7a5* expression levels in neural cells perinatally (Figure 2A-B). This time course again highlights a temporal dependence of *Slc7a5* function in postnatal neurodevelopment. Considering that one potential explanation for the observed phenotype is an excessive neural cell death in the postnatal period, we assessed the protein level of Cleaved (Cl)-Caspase-3, a main pro-apoptotic marker, on cortical samples obtained from control and mutant mice at different time points of development. Indeed, compared to controls, mutant animals show a significant increase in Cl-Caspase-3, specifically at P2 and P5 (Figure 4D), again supporting a role of *Slc7a5* during this time window. Importantly, the period affected by this surge in Cl-Caspase-3 levels corresponds to the phase of programmed cell death usually targeting cortical excitatory neurons. This is an innate process required to refine the number of neurons in the cerebral cortex (Blanquie et al., 2017). Specifically, to obtain a calibrated network, cortical neurons are generated in excess and subsequently eliminated by at least two waves of apoptosis, one of which occurs early after birth. This wave of programmed cell death represents a means to refine the cortical circuit after the phase of massive proliferation characterizing earlier developmental stages. In addition, previous studies have shown that to obtain an optimal pyramidal/inhibitory neuron ratio, the wave of programmed cell death affecting cortical excitatory neurons around P5 is followed by an adjustment in the number of inhibitory neurons (Wong et al., 2018). Thus, despite that *Slc7a5*^fl/fl^;*Emx1-*Cre+ mice only lack *Slc7a5* expression in the excitatory neurons of the forebrain, we thought that inhibitory cell number might be indirectly affected. Indeed, we found that, compared to their littermate controls, adult *Slc7a5*^fl/fl^;*Emx1*-Cre+ animals have a significantly lower number of inhibitory neurons, particularly in the upper cortical layers (Figure S6G-H), those layers directly affected by *Slc7a5* mutations (Figure S6D-F’). In contrast, the number of non-neuronal cells is entirely unaffected (Figure S6I-L), indicating that *Slc7a5* is important specifically for neuronal functions.

**Figure 4.**
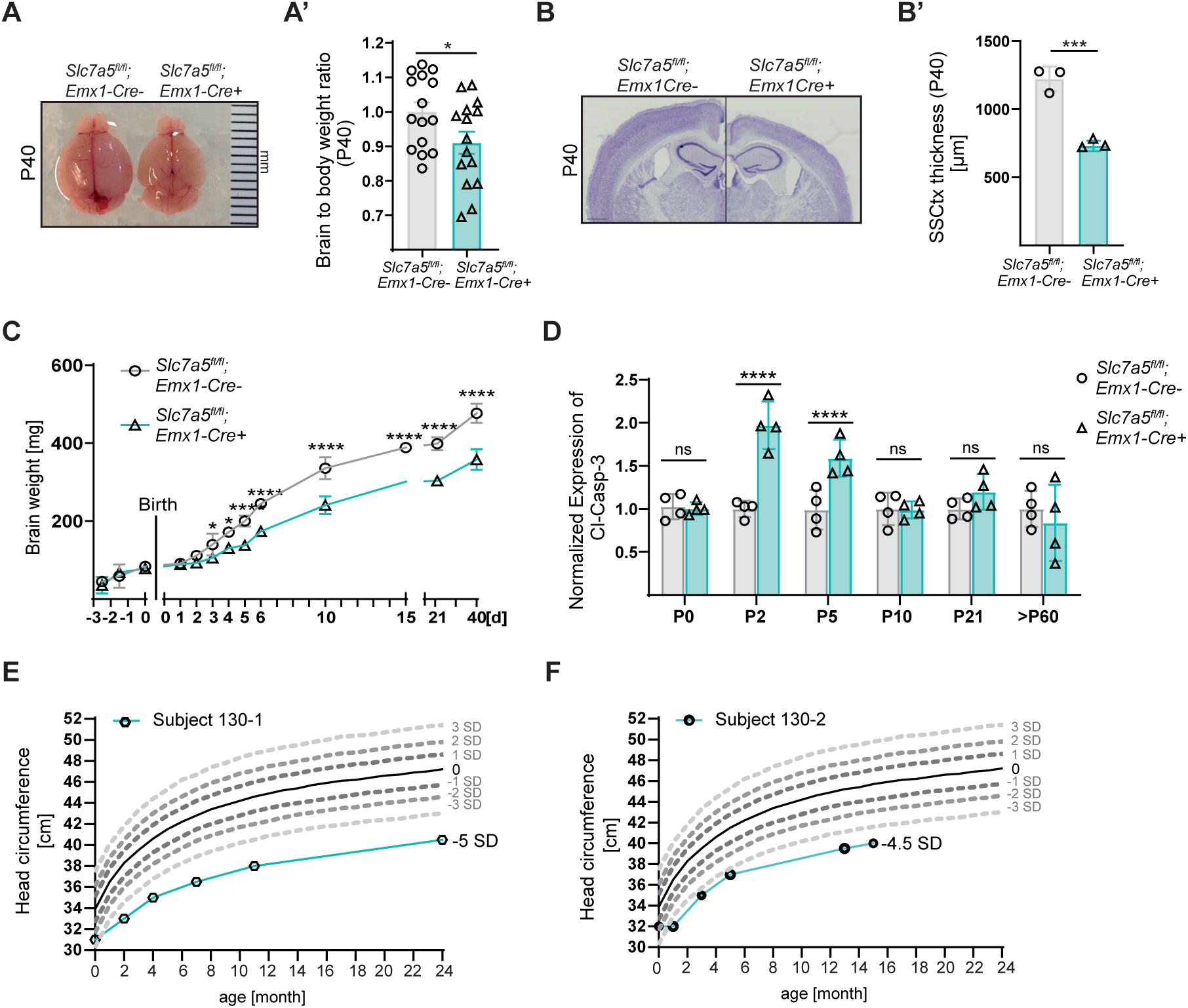
*Slc7a5* deficiency causes postnatal microcephaly. (**A**) Adult *Slc7a5^fl/fl^;Emx1-Cre+* mice present with reduced brain sizes. (**A’**) Quantification of brain to body weight ratio of adult *Slc7a5^fl/fl^;Emx1-Cre+* mice and littermate controls (means ± SEM; n = 15 mice per genotype; *\*p < 0.05*; unpaired two-tailed *t*-test). (**B,B’**) Nissl-staining of coronal mutant and wild-type brain sections show a reduction in cortical thickness in adult *Slc7a5^fl/fl^;Emx1-Cre+* animals. (**B**) Representative image of mutant and control coronal sections at P40, scale bar: 1500μm; (**B’**) Quantification of cortical thickness (means ± SEM; n = 3 littermates per genotype; *\*\*\*p < 0.001*; unpaired two-tailed *t*-test). (**C**) *Slc7a5^fl/fl^;Emx1-Cre+* mice develop postnatal microcephaly during the first 10 days after birth (means ± SEM; n ≥ 3 littermates per genotype and time point; *\*p < 0.05; ***p < 0.001*; *\*\*\*\*p < 0.0001;* multiple unpaired two-tailed *t*-test). (**D**) Western blot analysis of Gapdh-normalized expression of the Cleaved-Caspase-3 in mutant and wild-type cortical lysates at different developmental time points (means ± SEM; n = 4 mice per genotype and time point; *^ns^p>0.05; ****p < 0.00001*; multiple unpaired two-tailed *t*-test). (**E,F**) Measurements of the head circumference of patient 130-1 and 130-2 showing the progression of microcephaly over the first few months of life.

Next, we asked if the critical temporal window identified in mice is also sensitive to loss of *SLC7A5* function in humans. Thus, we identified new patients with mutations in *SLC7A5* and measured their head size for several weeks starting immediately after birth (Figure 4E-H). Patients 130-1 and 130-2 are two siblings from a non-consanguineous family presenting with the clinical feature of *SLC7A5* deficiency (Figure S6M). Extended Trio-WES analysis of both siblings and their parents identified compound heterozygous pathogenic variants in the *SLC7A5* gene: a previously described and functionally assessed missense variant c.1124C>T, p.(Pro375Leu) (Tărlungeanu et al., 2016) as well as a novel intragenic deletion of exons 5 to 10 with an assumed loss-of-function consequence. Both parents are heterozygous carriers of one *SLC7A5* pathogenic variant. Patient 130-1 showed microcephaly at birth (head circumference (HC) of 31 cm, -3 SD). The microcephaly progressively worsened to -5 SD (HC of 36.6 cm) at the age of 7 months (Figure 4F). A premature closure of fontanelles was observed from the age of 8 months. The history and presentation of the younger sibling, patient 130-2, are essentially similar. A mild microcephaly was diagnosed at birth (-2,5 SD), progressively deteriorating to -4.5 SD at the age of 6-7 months and followed by premature closure of fontanelles (Figure 4H). The phenotypic comparison of published patients (Tărlungeanu et al., 2016) revealed that the constant features associated with biallelic pathogenic *SLC7A5* variants include primary microcephaly, global developmental delay, seizures, autistic features and motor delay (Table S1). Thus, we concluded that *SLC7A5* mutations lead to microcephaly during the period of cortical refinement and programmed cell death in mice and humans.

### Pyramidal neuron loss is cell-autonomously linked to *Slc7a5* deficiency

Next, we investigated whether *Slc7a5* deficiency-linked neuronal cell death is due to a cell-autonomous or non-cell-autonomous effect. This was important especially in light of the observation that in the absence of Slc7a5 amino acids accumulate in the extracellular space, which could be potentially harmful to the tissue. Thus, to perform a quantitative assessment, we made use of the mosaic analysis with double markers (MADM) system, which facilitates concurrent fluorescent labeling and gene knockout in sparse single-cell clones *in vivo* (Contreras et al., 2021; Zong et al., 2005). Specifically, two reciprocally chimeric marker genes (*GT* and *TG* alleles) are targeted to identical loci upstream of *Slc7a5* (Chr8). The marker genes are part of the so-called MADM cassette (M8), which consists of split coding sequences for green (eGFP) and red (tdTomato) fluorescent proteins interspaced by a *loxP* site (Figure S7). Following Cre recombinase-mediated interchromosomal recombination, the sequence for eGFP and tdTomato are reconstituted. Due to an innately low stochastic interchromosomal recombination rate, the green (eGFP+), red (tdTomato+), and yellow (GFP+/tdTomato+) labeling are confined to individual sparse clones. In our experimental set-up, the Cre recombinase expression is coupled with the *Emx1* promotor. This facilitates MADM labeling and deletion of *Slc7a5* in single telencephalic radial glia progenitors (RGPs) and their progeny (Figure 5A), thereby generating cortex-specific genetic mosaics. To analyze potential cell-autonomous effects of the loss of *Slc7a5* in the developing neocortex, we assessed the relative abundance of green (eGFP, *Slc7a5*^-/-^) and red (dtTomato, *Slc7a5*^+/+^) excitatory neurons at different time points of postnatal development (P0, P5 and P40) in mosaic-MADM (*MADM-8^GT/TG,Slc7a5^;Emx1-Cre+*) and control-MADM (*MADM-8^GT/TG^;Emx1-Cre+*) animals (Figure 5 B-D’). While at P0 we did not observe significant changes in the ratio of green to red cells (Figure 5B-B’), indicating that Slc7a5 deficiency does not affect the proliferative phase of cortical development, by P5 there are significantly less *Slc7a5*^-/-^ than *Slc7a5*^+/+^ excitatory neurons (Figure 5C-C’). Mosaic-MADM animals present with significantly fewer *Slc7a5*^-/-^ (green) neurons in upper cortical layers (LII-LIV) compared to control (red) cells. In contrast, neurons in the lower layers (LV-VI) are not affected by *Slc7a5* deletion. The same analysis done at P40 revealed a slightly more profound reduction of mutant neuronal cell number in Mosaic-MADM (Figure 5D-D’). These findings are in accordance with the morphological changes observed in *Slc7a5*^fl/fl^;*Emx1-*Cre+ mice, where we observed a reduced thickness and cell density of the upper cortical layers (Figure S6D-F’). Altogether, this analysis indicates that sparse deletion of *Slc7a5* leads to a cell-autonomous increase in neuronal cell loss immediately after birth.

**Figure 5.**
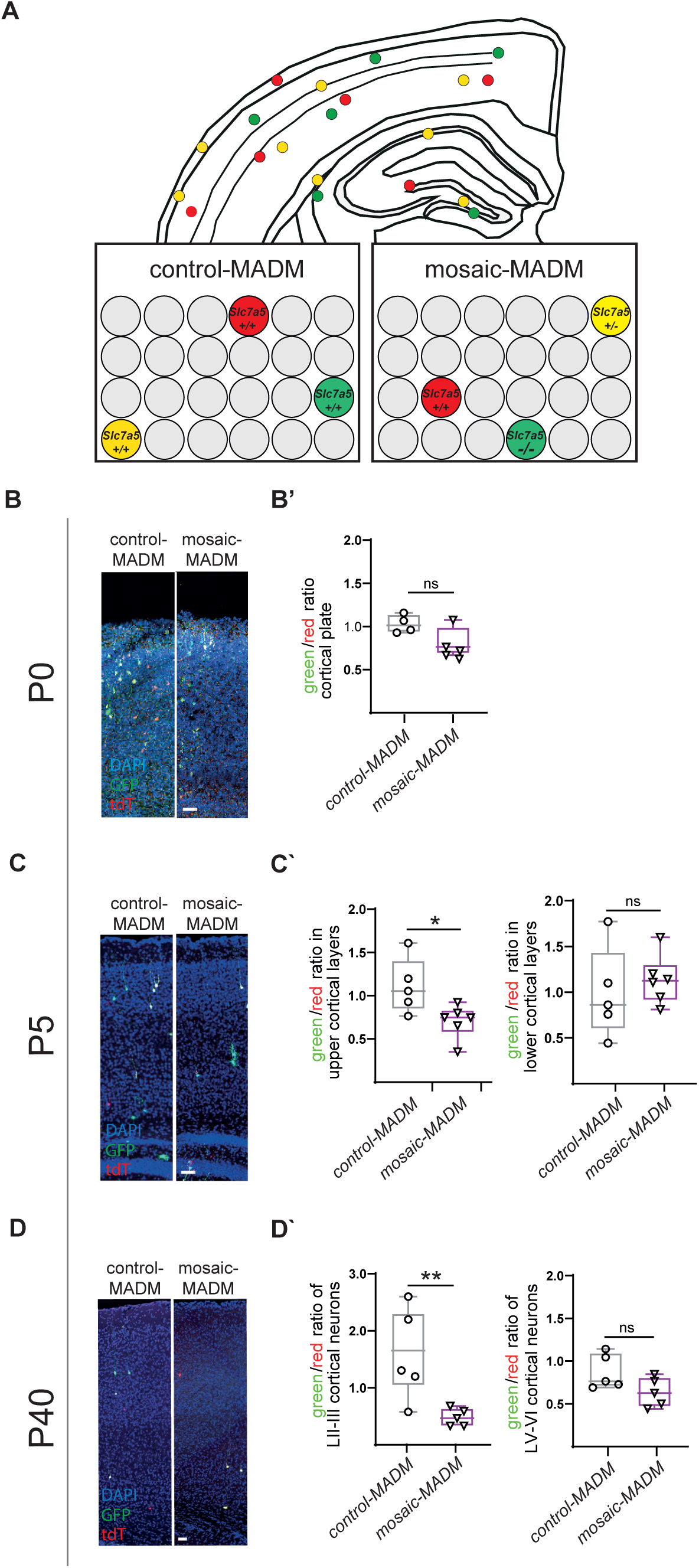
Loss of *Slc7a5* leads to a cell-autonomous reduction of cortical upper layer neurons. (**A**) Summary of the MADM principle: schematic representation and genotypes of cells of control-MADM (*MADM-8^GT/TG^;Emx1-Cre+)* and mosaic-MADM (*MADM-8^GT/TG,Slc7a5^;Emx1-Cre+)* cerebral cortex. (**B**) Coronal sections of P0 *MADM-8^GT/TG,Slc7a5^;Emx1-Cre+* and *MADM-8^GT/TG^;Emx1-Cre+* cortex and quantification (**B’**) of green/red cell ratio of MADM-labeled excitatory neurons in the cortical plate of mosaic-MADM animals at P0 compared to control-MADM littermates (n=4 animals per genotype; average of > 5 sections per animal; *^ns^p>0.05*; Mann-Whitney test; scale bar: 100μm). (**C**) Coronal sections of P5 *MADM-8^GT/TG,Slc7a5^;Emx1-Cre+* and *MADM-8^GT/TG^;Emx1-Cre+* cortex and quantification (**C’**) of green/red cell ratio of MADM-labeled upper layer (**left**) and lower layer (**right**) excitatory neurons in mosaic-MADM animals at P5 compared to control-MADM littermates (n=5 animals per genotype; average of > 5 sections per animal; *^ns^p>0.05; *p<0.05*; Mann-Whitney test; scale bar: 100μm). (**D**) Coronal sections of P40 *MADM-8^GT/TG,Slc7a5^;Emx1-Cre*+ and *MADM-8^GT/TG^;Emx1-Cre+*-cortex and quantification (**D’**) of green/red cell ratio of MADM-labeled upper layer (**left**) and lower layer (**right**) excitatory neurons in mosaic-MADM animals at P40 compared to control-MADM littermates controls (n=5 animals per genotype; average of > 5 slices per animal; *^ns^p>0.05;**p<0.001*; Mann-Whitney test; scale bar: 100μm).

### Large neutral amino acid-dependent metabolic reprogramming controls neuronal excitability in neonatal mice

What mechanisms can be underlying the stage- and cell-type-specific cellular phenotype observed in the *Slc7a5*-deficient cerebral cortex? We hypothesized that abnormal amino acid and lipid metabolism might ultimately lead to changes in neuronal excitability. Indeed, the role of lipids in modulating the activity of many ion channels is well-recognized (Elinder and Liin, 2017) and neuronal excitability is a factor determining the survival of cortical pyramidal cells during the postnatal wave of programmed cell death (Blanquie et al., 2017; Wong et al., 2018). Thus, we assessed the intrinsic excitability of layer II-III pyramidal neurons from the somatosensory cortex of mutant and control animals at P6-P7 by performing whole-cell current clamp recordings while applying a series of current steps to elicit action potentials (APs). Recordings from *Slc7a5^fl/fl^;Emx1-Cre+* mice and their wild-type littermates revealed a substantial reduction in neuronal firing in mutant animals (Figure 6A and Figure S8A). Given that in *Slc7a5^fl/fl^;Emx1-Cre+* mice both metabolic abnormalities and reduced neuronal survival are limited to the first days after birth, we performed current clamp recordings at P25-26 to assess whether the neuronal excitability defect is restricted to the same time window. Indeed, at P25-26 *Slc7a5^fl/fl^;Emx1-Cre+* samples are indistinguishable from controls (Figure 6B and Figure S8B), underscoring the importance of Slc7a5 in modulating neuronal excitability only early after birth. However, while, the transient nature of the phenotype suggests a rather direct link between the metabolic state of the neuron and its excitability, it remained a possibility that the observed electrophysiological abnormalities are due to plasticity effects associated with network properties. However, should the reduced excitability directly be connected to, or even cause Slc7a5-dependent neuronal cell loss, we expected this phenotype to be cell-autonomous. We therefore again took advantage of the mosaic-MADM mouse model (Figure 6C) and performed current clamp recordings from *Slc7a5*^-/-^ (green) and *Slc7a5*^+/+^ (red) excitatory neurons in the same mosaic animal. Similar to what we observed in *Slc7a5^fl/fl^*;*Emx1-Cre+* mutants, the AP firing rate triggered by current injection was significantly reduced in LII-III *Slc7a5*^-/-^ neurons from the somatosensory cortex at P6-7 (Figure 6D). Altogether, these analyses indicate that *Slc7a5* expression cell autonomously controls neuronal excitability specifically at early stages after birth. To understand the potential bases of the reduced firing rate caused by *Slc7a5* deletion, we performed a detailed analysis of the AP properties. We immediately inferred that *Slc7a5* mutant neurons at P6-7 do not fire less because they are more immature than wild-type neurons since they show AP features rather more comparable to those observed in older (i.e., P25-26) control neurons (Figure 6E). Indeed, compared to controls, mutant neurons at P6-7 display larger amplitude (Figure 6F), faster rise time (Figure 6G) and more hyperpolarized AP threshold (Figure 6H), matching the AP features observed in control neurons at P25-26 and suggesting a different modulation of the voltage-gated channels involved in the initiation and rising phase of APs. On the other hand, other properties such as the resting membrane potential, the inter-spike interval (ISI) ratio (calculated as the ratio of the last ISI relative to the first), the AP decay time and half-width are unchanged at P6-7 (Figure S8C-F). We further performed a closer examination of the AP waveform using the phase plane plot analysis (Figure 6I) to evaluate the dynamic changes of the membrane potential over time (dV/dt; Figure 6J) and to better separate the different phases of the AP. Our analysis revealed a striking increase in the velocity of the AP, highlighting a potential defect at the axon initial segment (AIS) and a faster AP backpropagation in the somatodendritic compartment of mutant neurons (Figure 6I-J). While the unchanged resting membrane potential would exclude a role of ATP-dependent potassium channels, these analyses point to changes in the properties of multiple channels, probably through a different modulation and/or distribution of voltage-gated sodium (e.g., Nav1.2) and potassium channels (e.g., A-type).

**Figure 6.**
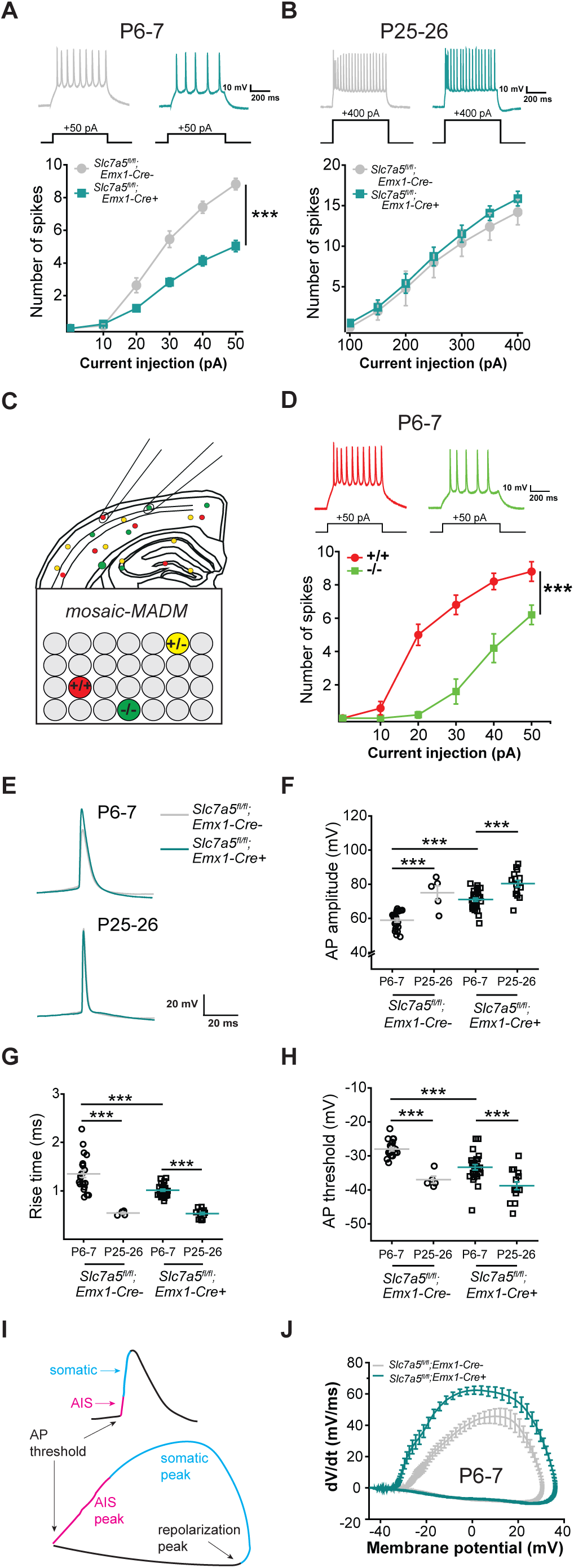
Intracellular amino acid levels modulate neuronal excitability perinatally. (**A**) Current clamp recordings from LII/III pyramidal neurons in *Slc7a5^fl/fl^;Emx1-Cre+* and *Slc7a5^fl/fl^;Emx1-Cre-* somatosensory cortex (SSCtx) at P6-P7 (*Emx1Cre*-: n = 22 cells / 3 mice; *Emx1-Cre*+: n = 30 cells / 3 mice; Two-way ANOVA: genotype F(1,311) = 123.01 ***p<0.001, current step F(5,311) = 205.3 ***p<0.001, interaction F(5,311) = 16.75 ***p<0.001) and (**B**) P25-P26 (*Emx1Cre*-: n = 5 cells / 3 mice; *Emx1-Cre*+: n = 15 cells / 3 mice; Two-way ANOVA: genotype F(1,139) = 1.84 ^ns^p>0.5, current step F(6,139) = 34.87 ^ns^p>0.5, interaction F(6,139) = 0.07 ^ns^p>0.5). (**C**) Studying the cell-autonomous nature of the reduced firing rate by using whole cell patch clamp recordings from green (Slc7a5-/-) and red (Slc7a5 +/+) LII/III pyramidal neurons from the same mosaic-MADM animal. (**D**) Current clamp recordings from LII/III pyramidal neurons in SSCtx of P6-P7 mosaic-MADM mice (Slc7a5 +/+: n = 5 cells / 5 mice; Slc7a5 -/-: n = 7 cells / 5 mice; Two-way ANOVA: genotype F(1,65) = 116.05 ***p<0.001, current step F(5,65) = 79.22 ***p<0.001, interaction F(5,65) = 11.16 ***p<0.001). (**E**) Representative AP traces from current clamp recordings of data shown in (**A-B**). The action potential (AP) amplitude (**F**), AP rise time (**G**) and AP threshold (**H**) are transiently affected in Slc7a5 deficient LII/III pyramidal neurons at P6-7 (*Emx1-Cre*-: n = 22 cells / 3 mice; *Emx1-Cre*+: n = 30 cells / 3 mice) compared to age-matched littermates and P25-26 time point (*Emx1-Cre*-: n = 5 cells / 3 mice; *Emx1-Cre*+: n = 15 cells / 3 mice). Two-way ANOVA for AP amplitude: genotype F(1,71) = 23.15 ***p<0.001, time point F(1,71) = 52.57 ***p<0.001, interaction F(1,71) = 3.24 ^ns^p>0.5, Holm-Sidak post hoc ***p<0.001. Two-way ANOVA for AP rise time: genotype F(1,71) = 7.32 ***p<0.001, time point F(1,71) = 101.21 ***p<0.001, interaction F(1,71) = 6.4 *p<0.05, Holm-Sidak post hoc ***p<0.001. Two-way ANOVA for AP threshold: genotype F(1,71) = 10.56 **p<0.01, time point F(1,71) = 42.93 ***p<0.001, interaction F(1,71) = 2.61 ^ns^p>0.5, Holm-Sidak post hoc ***p<0.001. (**H**) Representative AP traces recorded from LII/III pyramidal neurons in *Slc7a5^fl/fl^;Emx1-Cre+* and *Slc7a5^fl/fl^;Emx1-Cre-* SSCtx at P6-7 and P25-26. (**I**) AP plotted as voltage vs. time (**top**) and dV/dt vs. voltage (phase-plane plot, **bottom**). Different phases of the AP are color-coded across panels to highlight different phases of the AP corresponding to initiation of AP in axon initial segment (AIS; pink), propagation in the somatodendritic compartment (blue), and the repolarization peak. (**J)** Phase-plane plot of data shown in (**A**) reveals defects in the AIS and the somatodendritic compartment in LII/III pyramidal neurons of *Slc7a5^fl/fl^;Emx1-Cre+* animals.

### *Slc7a5* deficient animals show persistent behavioral defects

To assess whether the alterations observed in *Slc7a5* mutant animals during the period of cortical circuit refinement lead to permanent behavioral abnormalities, we subjected adult *Slc7a5^fl/fl^;Emx1-Cre+* and control animals to different behavioral tests. We found that, in an open field, mutant animals present with increased horizontal (Figure 7A-B’) and vertical explorative behavior (Figure 7C), indicative of a hyperactivity phenotype. However, in the same setting *Slc7a5^fl/fl^;Emx1-Cre+* animals did not display differences in anxiety-like behaviors (Figure 7D). Since *SLC7A5* patients present with severe motor deficits, we assessed locomotion features in *Slc7a5*^fl/fl^*;Emx1-Cre+* mice. Indeed, mutant mice exhibit moderate motor defects indicated by gait abnormalities (Figure 7E), such as decreased stride and stance length (Figure 7F-F’’), as well as hind limb clasping behavior (Figure 7G-G’). Interestingly, since the *Emx1* promoter is not active in the cerebellum, the locomotion abnormalities observed in the Slc7a5^fl/fl^*;Emx1-Cre+* mouse model have a forebrain origin. Next, using the three-chamber sociability test, we tested *Slc7a5^fl/fl^;Emx1-Cre+* mice for social interest and social memory behaviors. Wild-type animals commonly prefer interacting with a stranger mouse (M) over an object (O) or with a novel stranger mouse (M2) over the familiar mouse (M1). In mutant mice, these preferences are not detectable (Figure 7H-K’), indicative of sociability and social-memory problems. In summary, our analyses suggest that, although the direct effects of *Slc7a5* deficiency (i.e., decreased intake of essential neutral amino acids) are limited to a specific and very short time window, *Slc7a5* mutations cause persistent cortical-circuit and behavioral dysfunctions.

**Figure 7.**
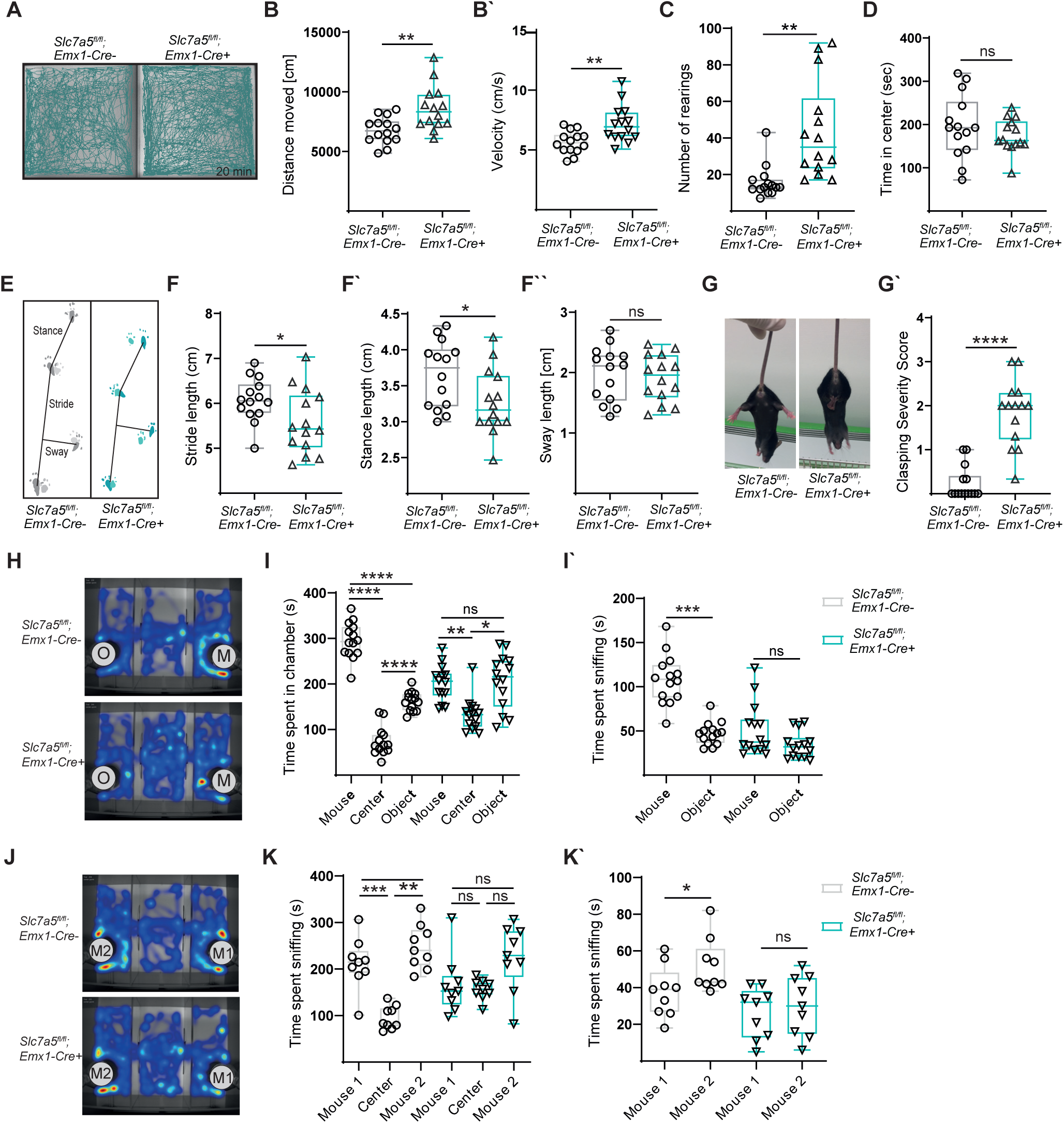
Loss of *Slc7a5* in cortical excitatory neurons causes persistent behavioral dysfunctions. (**A-C**) *Slc7a5^fl/fl^;Emx1-Cre+* animals present with a hyperactivity phenotype. (**A**) Cumulative traces of mutant and wild-type animals of a 20 min recording session in the open field. (**B**) Quantification of the total distance traveled, (**B’**) velocity, (**C**) number of rearings and (**D**) time spent in center during one open field session (n=14 mice per genotype; two-tailed unpaired *t*-test). (**E-F’’**) *Slc7a5^fl/fl^;Emx1-Cre+* animals display a mild gait defect. (**E**) Representative control and mutant strides. Quantification of stride (**F**), stance (**F’**) and sway (**F**’’) length (n=14 per genotype, *\*p < 0.05*; two-tailed unpaired *t*-test). (**G**) Hind limb clasping observed in adult *Slc7a5^fl/fl^;Emx1-Cre+* mice. **(G‘)** Scoring of hind limb clasping severity (0–1 (normal) to 3 (most severe) (n = 14 animals per genotype; *\*\*\*\*p < 0.00001*; two-tailed unpaired *t*-test). (**H-K’**) *Slc7a5^fl/fl^;Emx1-Cre+* mice present mild defects in sociability in the three-chamber social interaction test (TCST). (**H**) Representative heat maps of control (top) and mutant (bottom) behavior during the first round of the TCST. (**I**) Quantification of time spent in chamber and (**I‘**) time spent sniffing (*n* = 14 mice per genotype; *\*\*\*\*p < 0.00001*, *\*\*p < 0.01*; *\*p < 0.05; ^ns^p > 0.05*; 1-way ANOVA and Sidak’s multiple comparison test). (**J**) Representative heat maps of control (top) and mutant (bottom) behavior during the second round of the TCST. (**K**) Quantification of time spent in chamber and (**K‘**) time spent sniffing (*n* = 9 mice per genotype; females only; *\*\*\*p < 0.0001*, *\*\*p < 0.01*; *\*p < 0.05; ^ns^p > 0.05*; 1-way ANOVA and Sidak’s multiple comparison test).

## DISCUSSION

Neurons are generated in large amounts early during embryonic brain development but a significant fraction of them are removed at subsequent developmental stages (Dekkers et al., 2013; Nikolić et al., 2013; Southwell et al., 2012). The removal of these cells must be highly selective and therefore regulated by very tight mechanisms, possibly integrating both extrinsically- and intrinsically-driven processes. While a complete view of the factors directing this process is still missing, the literature suggests that neuronal activity might be used as a measure of neuronal participation in the network, and therefore it is a central determinant of the programmed cell death-based refinement of the perinatal network (Dekkers et al., 2013; Wong et al., 2018). However, the nature of potential upstream signaling and the pattern of neuronal activity determining this phenomenon remain unclear. Identifying extrinsic and intrinsic factors that can modulate neuronal properties at this perinatal developmental stage is critical since affecting this process, even shortly, can permanently affect brain circuits. Here we focused on the metabolic program of neural cells of the cerebral cortex as a measure of the intrinsic fit of a neuron and a determinant of its integration in the cortical circuit. Although metabolism is a crucial determinant of cellular fitness and neurons might be exposed to different metabolic environments, a description of how the level of various metabolites changes in the forebrain over time was still missing. By obtaining a metabolome profile of the cortex at various developmental stages, we provide a comprehensive view of the metabolites detected in this brain region and their changes over the course of development. As several metabolites are linked to neurodevelopmental conditions, our data also represents an important starting point to evaluate potential critical time windows in the context of brain disorders. For example, our analysis underscored a downward trajectory for essential large neutral amino acids, with their level becoming significantly lower in the cerebral cortex at the perinatal period. By deleting *SLC7A5*, a large neutral amino acid transporter whose mutations have been recently identified as a cause of autism and microcephaly, we tested the importance of regulating large neutral amino acid levels for the metabolic and physiological state of neural cells. We found that *SLC7A5* expression is a determinant factor in specifying cortical neurons’ metabolic state, particularly at perinatal stages. In this context, it is very intriguing to observe that *SLC7A5* transcription in neurons is induced by hypoxia (Fitzgerald et al., 2021; Onishi et al., 2019), a physiological state fetuses experience during and shortly after birth (Huch et al., 1977). What happens if the typical level of Slc7a5 substrates is not met during this developmental window? We report that a decreased level of branched-chain amino acids is coupled with a disturbance of lipid metabolism and neuronal excitability, providing an elegant example of coupling the intrinsic fitness of a cell with its integration in the neuronal network. Our mosaic analysis further suggests that intrinsic excitability can directly affect neuronal survival probability at this developmental stage. The exact mechanisms underlying the reduced neuronal excitability remain unclear. Our transcriptomic and proteomic analysis did not uncover changes in ion channel expression in *Slc7a5* deficient neurons, indicating that the change in neuronal excitability must be due to another regulatory mechanism. The most plausible explanation is that the shift in lipid profile observed in mutant cells in neonatal mice leads to a different modulation of some ion channels involved in neuronal excitability. Altogether, our analysis highlights the importance of dietary-obtained factors, such as essential amino acids, for neurodevelopment. Importantly, the similar trajectory of the microcephaly onset observed in mice and humans with *SLC7A5* mutations suggests that although our metabolic profile describes changes in the murine brain, humans and mice may employ a similar metabolic program across time. Furthermore, the stage and cell-type specificity of the effects observed in *SLC7A5* mutants points to the importance of performing longitudinal studies evaluating environmental, metabolically-relevant factors that can influence particular stages of brain development and that may interact with genetic factors associated with human neurodevelopmental conditions.

## METHODS

### Mice

The *Slc7a5*^fl^ (B6.129P2-Slc7a5tm1.1Daca/J) mouse line in which the loxP sites are flanking exon 1, including the initiation codon, was used. The *Slc7a5^fl^; Emx1-Cre* conditional line was generated by crossing *Slc7a5^fl/fl^* with *Slc7a5^+/fl^* animals expressing Cre recombinase under the *Emx1* promotor (B6.129S2 (Emx1tm1cre)Krj/J). *Slc7a5^fl^;Tie2-Cre* conditional animals were generated by crossing *Slc7a5^fl/fl^* Cre-negative females with *Slc7a5^+/fl^*;*Tie2-Cre* positive males. Genotyping for the loxP site was performed using the following primers: forward CCA TCT CGG CAG TTC CAG GC and reverse GGT GCT TTG CTG AAG GCA GGG. Potential germline recombination events were closely monitored using delta-primers: forward CAG CTC CTT TCT CCA GTT AAG C and reverse GAC AGC CTG AAG TAA AAT TCC C. Presence of Cre recombinase was assessed using the primers: forward GTC CAA TTT ACT GAC CGT ACA CC and reverse GTT ATT CGG ATC ATC AGC TAC ACC. Generation of the *Tie2-Cre*, *Emx1-Cre* and *Slc7a5*^fl^ lines has been described previously (Gorski et al., 2002; Kisanuki et al., 2001; Sinclair et al., 2013). We generated mosaic *Slc7a5*-MADM (*MADM-8^GT/TG,Slc7a5^;Emx1^-Cre/+^*) mice with sparse green GFP^+^ homozygous *Slc7a5^-/-^* mutant cells, yellow (GFP^+^/tdTomato^+^) heterozygous *Slc7a5^+/-^*, and red (*tdTomato^+^*) *Slc7a5^+/+^* wild-type cells in an otherwise unlabeled heterozygous background. To this end, *Slc7a5* floxed alleles were genetically linked to the MADM-8 TG cassette via meiotic recombination as described previously (Contreras et al., 2021). Genotyping for the presence of loxP sites and Cre recombinase was performed as described above. A detailed description of the primer sequences for MADM-GT and MADM-TG cassettes for MADM-8 can be found in Contreras et al. 2021. Embryonic time points were determined by plug checks after timed matings, defining embryonic day (E) 0.5 as the morning after copulation. Animals were kept on a 12 hour light/dark cycle (lights on at 7:00 am) and housed in groups of 3-4 animals per cage, with food and water available ad libitum. Experiments were carried out under specific pathogen-free conditions and the health status of the mouse line was routinely checked by a veterinary. Experiments were carried out with littermates of the same sex. Both males and females were used for experiments. All animal protocols were approved by the Institutional Animal Care and Use Committee at IST Austria and the Bundesministerium für Bildung, Wissenschaft und Forschung, Austria, approval number: BMBWF-66.018/0015-V/3b/2019.

### Human Patients

Family 130 with two affected daughters (patients 130-1 and 130-2) was referred to genetic counseling and testing in the Institute of Medical Genetics of Medical University Vienna through a supporting Centre for Developmental Disabilities in Vienna. A detailed natural history of the development of the phenotype, notably of microcephaly is available in both siblings. Patient 130-1 was born at term and showed microcephaly at birth (head circumference (HC) of 31 cm, -3 SD). Birth weight and height were normal. The microcephaly deteriorated progressively to -5 SD (HC of 36.6 cm) at the age of 7 months. A premature closure of fontanelles was observed from the age of 8 months and a surgical treatment (frontobasal advancement) of presumed craniosynostosis was performed at the age of 12 months. The patient displayed pronounced muscular hypotonia and motor delay. At the age of 3 years, seizures started that could be successfully treated with valproate. In addition to microcephaly, brain MRI also showed pontocerebellar and corpus callosum hypoplasia. At the time of first genetic counseling, the patient 130-1 was 9 years and 4 months old and presented with severe microcephaly as well as a growth retardation (HC 43 cm, -7 SD; height 109 cm, -4.5 SD and weight 17.1 kg, -4 SD). She could sit independently and stand with help, however not walk. The speech was absent. The history and presentation of the younger sibling, patient 130-2, are essentially similar. The microcephaly was diagnosed at birth, progressively deteriorating to -4.5 SD at the age of 6-7 months with following premature closure of fontanelles. Pontocerebellar and corpus callosum hypoplasia were detectable by brain imaging. At genetic counseling, the patient 130-2 was 4 years and 4 months old and displayed severe microcephaly (-6 SD), growth retardation (-5 up to -6 SD), global developmental delay (independent sitting possible, no walking, absent speech) and autistic features. The first seizures occurred at the age of 6 years and are well controlled with anticonvulsive treatment. Prior to WES analysis an aCGH with normal results was carried out in both patients. <u>Whole-exome sequencing of patient samples:</u> We performed whole-exome sequencing in both affected siblings and their healthy unrelated parents (extended Trio-WES). Written informed consents were obtained from the whole family involved. The DNA samples were prepared following the workflow of the TruSeq Exome Library Kit (Illumina) for the enrichment of exonic regions. The final library was paired-end sequenced on an Illumina NextSeq500 sequencer. The obtained sequencing reads were aligned to GRCh37/hg19 using the Burrows Wheeler Aligner (BWA-MEM) and further processed in house according to GATKs best practice protocol for calling single nucleotide variants, insertions and deletions. The evaluation of the called variants was performed using the program VarSeq from Golden Helix Inc®. Variants were classified according to the American College of Medical Genetics and Genomics (ACMG) guidelines (Richards et al., 2015). In addition, a copy number variation (CNV) analysis was performed for all analyzed samples comparing the calculated coverage of each sequenced sample to the already existing coverage data, obtained from BAM-files, for all previously analyzed in-house samples. This analysis was also done by a supported module from VarSeq within the VarSeq software from Golden Helix®. <u>Quantitative PCR (qPCR) of patient samples:</u> To verify the identified multiexonic deletion of *SLC7A5* we performed a quantitative PCR (qPCR) using CFX Connect^TM^ Real-Time PCR Detection System (Bio-Rad) with primers spanning the deleted region. All primers were purchased from Eurofins Genomics. The specificity of each primer set was monitored by a dissociation curve. PCR reactions were performed in triplicate and normalized to *PAPD5 and PRKD1*. Primer sequences are available on request.

### Immunofluorescence staining

<u>Immunofluorescent staining in adult mice:</u> Adult littermate animals were anesthetized and transcardially perfused with 4% paraformaldehyde (PFA). After dissection, brains were postfixed in 4% PFA and dehydrated using 30% sucrose (in 1X PBS). Dehydrated brains were sectioned at 30-40 μm on a Leica Sliding Microtome SM2010R. Adult brain sections were stained free-floating. No antigen retrieval was performed unless specifically recommended in the primary antibody datasheet. Sections were washed in 1x PBS and blocked with 4% normal donkey or goat serum. Primary antibodies were diluted in 0.3% Triton X-100 and 4% serum and incubated overnight on a horizontal shaker at 4°C. After 14-16h, the sections were washed and a species-specific secondary antibody was added for 2 hours. 300 nM DAPI (Life Technologies) in 1x PBS was applied for 10min to achieve a nuclear counterstaining, followed by mounting in DAKO fluorescent mounting medium. <u>Immunofluorescent staining in embryos and pups</u>: In brief, embryos were quickly decapitated and the whole head was immersion-fixed overnight in 4% PFA at 4°C. Pups (>P0) were decapitated and the brain was dissected. After dissection the brain was immersion-fixed overnight in 4% PFA at 4°C. After fixation the samples were dehydrated in 30% sucrose and embedded in OCT Cryoprotectant. Brains were sliced on a Thermo Scientific CryoStar NX70 cryostat and directly mounted on Superfrost® microscope slides. Immunofluorescence stainings were performed directly on the slides in a humidity chamber. The detailed staining procedure is stated in the section above. <u>Quantification:</u> To examine cortical layering, the thickness of Cux1-positive cell layer and the Ctip2-positive cell layer was measured at three defined points of each cortical hemisphere (n= 3 littermate animals per genotype, at least 4 images/animal). The density of upper or lower layers neurons was quantified by normalizing the number of Ctip2+ or Cux1+ cells to the total amount of DAPI+ cells within a defined square. To assess the number of apoptotic cells, Cl-Caspase-3+ cells were quantified in coronal brain sections. The number of apoptotic cells in the cortex was normalized to the number of Cl-Casp3 positive cells in the rest of the brain (n= 3 littermate animals per genotype, at least 5 images/animal). To monitor a potential decrease in the number of inhibitory neurons, the number Parvalbumin positive cells was quantified per layer and normalized to the absolute number of PV+ cells per counting window (n= 3 littermate animals per genotype, at least 5 images/animal). The numbers of astrocytes and microglia were monitored by assessing the number of Gfap+ or Iba1+ cells per cortical column. The numbers were normalized to the size of the counting square. (n= 3 littermate animals per genotype, at least 5 images/animal).

The following primary antibodies were employed:

Anti-Ctip2 (ab18465) - rat - 1:500

Anti-Cux1 (sc-13024) - rabbit - 1:500

Anti-Iba1 (Wako 019 19741) - rabbit - 1:500

Anti-Gfap (CST 12389S) - rabbit - 1:500

Anti-Parvalbumin (ab11427) - rabbit - 1:500

<u>Imaging:</u> Images from immunofluorescent stainings were acquired on a Zeiss LSM800 inverted confocal microscope and analyzed in Fiji.

### Immunohistochemistry stainings - Nissl staining

<u>Nissl staining:</u> For Nissl staining, adult animals were transcardially perfused with 4% PFA and post-fixed overnight in 4% PFA at 4°C. P0-1 brains were immersion fixed in 4% PFA overnight at 4°C followed by a dehydration step with 30% sucrose (in 1X PBS). P0-1 coronal sections were obtained at the Thermo Scientific CryoStar NX70 cryostat. Adult coronal sections at 40 μm thickness were cut on a Leica Sliding Microtome SM2010R. Sections were mounted on Superfrost® microscope slides. Nissl staining was performed with 1% Cresyl Violet solution (Cresyl Violet Acetate, Sigma, Cat.No C 5042). Sections were cleared with RotiHistol (Carl Roth) for 10 min and rehydrated (absolute EtOH to water: 96%, 90%, 70%, 50%, 30%, water, 3-5 min each). Images were acquired with an Olympus Slide scanner VS120 and quantified using Fiji (n=3/genotype, at least 4 images/animal).

### Isolation and quantification of MADM-labelled tissue

<u>Tissue harvesting:</u> P5 and P0 animals were decapitated, brain was isolated and fixed in 4% PFA overnight. From P6 onwards, mice were deeply anesthetized by injection of a ketamine/ xylazine/ acepromazine solution (65 mg, 13 mg and 2 mg/kg body weight) and unresponsiveness was confirmed through pinching the paw. The diaphragm of the mouse was opened from the abdominal side to expose the heart. Cardiac perfusion was performed with ice-cold PBS (phosphate-buffered saline) followed immediately by 4% PFA (paraformaldehyde) prepared in PB buffer. Brains were removed and further fixed in 4% PFA O/N to ensure complete fixation. Brains were cryopreserved with 30% sucrose (Sigma-Aldrich) solution in PBS for approximately 48 h and were then embedded in Tissue-Tek O.C.T. (Sakura). Tissue was stored at -80°C until further usage. 40μm coronal frozen sections were cut on a Leica Sliding Microtome SM2010R, washed in 1X PBS and mounted on Superfrost glass slides. Next, sections were stained with 300 nM DAPI (Life Technologies) and mounted using Dako mounting medium. <u>Imaging:</u> Images were acquired using a Zeiss LSM800 inverted confocal, Olympus Slidescanner VS120 or Nikon Eclipse Ti2. Images were post-processed and analyzed in Zen Blue 2.6 software and Fiji. <u>Quantification:</u> The number of MADM-labelled green, red and yellow neurons within the somatosensory cortex was quantified according to their localization within the different cortical layers. The ratio between green and red neurons was calculated per animal (>10 hemispheres per animal; n=6 littermate pairs). A Mann-Whitney test was used for statistical analysis.

### Electron microscopy

<u>Sample preparation:</u> P2 *Slc7a5^fl^;Emx1-Cre* littermates (n=4 per genotype) were decapitated and dissected. The brains were dropped into EM suitable fixative (2.5% glutaraldehyde and 2% PFA in 0.1M PB) and fixed for 10 min using a Pelco BioWave Pro+ microwave. Brains were washed in 1X PB and sliced at a Leica vibratome (100μm thick slices). Slices were placed in 1% osmium tetroxide in 0.1M PB, followed by 1% uranyl acetate in 50% ethanol and a dehydration train with ascending ethanol concentration (50%, 70&, 90%, 96%, 100%, 100%). Samples were then placed into propylene oxide followed by 1:2, 1:1 and 2:1 Durcupan:propylene oxide and an overnight incubation in Durucupan. Samples were mounted on siliconized coverslips, placed on a heating plate for 30 min at 37°C and put in an oven for 2 days at 60°C to polymerize resin. The region of interest (layer 2/3 somatosensory cortex) was cut and re-embedded in a resin block for further slicing. 70 nm ultrathin serial sections were cut using an UC7 ultramicrotom (Leica Microsytems), collected on formvar-coated copper-slot grids and examined in FE-SEM. <u>Imaging:</u> Images were acquired on a Tecnai T10 transmission electron microscope at 24000X magnification.

### Western Blot

<u>Sample preparation:</u> Littermate *Slc7a5^fl^;Emx1-Cre* animals at different developmental time points (minimum n= 3 per genotype) were decapitated, the brain was dissected on ice, snap frozen and stored at -80°C. For the LAT1 expression time course in neural cells only *Slc7a5^fl/fl^;Tie2-Cre+* cortices at different developmental time points (E16.5- P60 n=1 per timepoint) were collected. For protein extraction cortices were homogenized in ice-cold lysis buffer (20mM Tris-HCl pH:8, 137mM NaCl. 10% Glycerol, 0.1% NP-40, 1mM EDTA, 9.5mM NaF, 10mM PPi (Sodium pyrophosphate dibasic), 1mM Na3Vo4) plus cOmplete™ Protease Inhibitor. Samples were kept for 20 min on ice followed by a centrifugation step at 500g for 10 min at 4°C. The supernatant was transferred into a fresh tube and centrifuged at max speed for 20 min at 4°C. The protein concentration of the lysates was determined using the Pierce™ BCA Protein Assay Kit (Thermo Fisher, Cat. no. 23225). <u>Western Blots</u>: Samples were diluted with 2X Laemmli buffer (10% SDS, 20% Glycerol, 100mM Tris-Cl (ph 6.8), Bromophenol blue, 3% DTT) and denatured at 65°C for 10 min and separated using 8-12% SDS-PAGE gels. In case of cortical lysates 25-50 μg of proteins were loaded per sample. Proteins were blotted to a PVDF membrane (Merck) for 1-2 h at 4°C with 300 mA constant current using the Bio-Rad immunoblot apparatus. The membranes were blocked with 2.5% BSA in 1x TBST for 1h at room temperature. Primary antibodies were diluted in blocking solution and incubated overnight at 4°C on a shaker. Secondary anti-IgG antibody coupled to horseradish peroxidase (HRP) was incubated for 1h at RT. For detection PierceTM enhanced chemiluminescent substrate (ThermoFisher) in combination with the GE Healthcare Amersham machine was used. <u>The following primary antibodies were employed:</u>

Anti Human L-type Amino Acid Transporter 1(LAT1) (KE026) - rabbit - 1:1000

Anti Cleaved-Caspase-3 (CST 9661) - rabbit - 1:1000

Anti-Gapdh (ABS16) - rabbit - 1:1000

Anti-S6 ribosomal protein (CST 2217) – rabbit - 1:1000

Anti-phosphoS6 ribosomal protein^240-244^ (CST 5364) – rabbit - 1:1000

Anti-phosphoS6 ribosomal protein^235-236^ (CST 2211s) – rabbit - 1:1000

Anti-4eibp1 (CST 9644) – rabbit - 1:500

Anti-phospho4ebp1 (CST 2855) – rabbit - 1:500

Anti-Atf4 (cat.no10835-I-AP) – rabbit - 1:500

Anti-Lamp1 (ab24170) - rabbit - 1:1000

Anti-Lc3-I/II (Millipore ABC929) – rabbit - 1:250

Anti-Eif2a (CST 9722) – rabbit - 1:500

Anti-phosphoEif2a (CST 9721S) – rabbit - 1:500

Anti-AMPK (CST 2532) – rabbit - 1:500

Anti-phosphoAMPK (CST 2535) – rabbit - 1:500

### Untargeted and targeted Metabolomic analysis of cortical tissue

<u>Sample preparation for untargeted metabolomic analysis of cortical tissue:</u> Cortices of *Slc7a5^fl/fl^;Emx1-Cre+ and Slc7a5^fl/fl^;Emx1-Cre-* animals were collected at three different developmental timepoints (E 14.5, P2 and P40) (n= 4 per genotype and time point). *Slc7a5^fl/fl^;Emx1-Cre-* animals are hereafter referred to as wild type. The tissue was collected in 2ml Eppendorf tubes, washed with 1ml ice-cold 1X PBS and weighed. The samples were stored at -80°C until further processed. For the extraction of metabolites the samples were thawed on ice. The brain tissue was combined with ice-cold solvent mixture (methanol:acetonitrile:H2O (2:2:1, v/v) MS-grade; cooled to – 20 °C). The brain tissue was homogenized with a Bel-Art® disposable pestles for 1 min followed by sonication for five minutes in a water bath sonicator at RT. The samples were incubated for 1 hour at -20 °C, followed by a centrifugation step at 14,000g for 3 min at 4°C. The supernatant was stored at -20°C until the next step and the pellet was re-suspended in ice-cold solvent (methanol:acetonitrile:H2O (2:2:1, v/v)). After vortexing the mixture for 1 min at 4°C, the samples were incubated for another hour at -20°C. After another centrifugation step at 14,000g for 3 min (4°C) the supernatant was combined with the previous supernatant and incubated for 2 hours at -20 °C. This was followed by a last centrifugation step at 14,000g for 10 min at 4°C. The supernatant was transferred to a new 1.5-ml microcentrifuge tube, snap-frozen and stored at -80°C.

<u>Sample preparation for intracellular targeted metabolomic analysis</u>: P2 *Slc7a5^fl^;Emx1-Cre* animals were decapitated (n=7/genotype). The brains were dissected on ice. Cortices of P2 *Slc7a5^fl^;Emx1-Cre* animals were dissected. Meninges and hippocampus were removed from the cortex. A single cell suspension was prepared according to the protocol provided by Papain Dissociation System - Worthington Biochemical Corp.. Cortices were moved to 50-ml TPP® TubeSpin Bioreactor tubes and further processed according to the kit’s manual. Cell pellets were washed and re- suspended in EBSS+1%BSA to avoid potential contamination with external metabolites. Single cell suspension was stored on ice in 1X EBSS+1%BSA while the live cell count/sample was determined using a TC20 Automated Cell Counter (Bio-Rad). For the extraction of metabolites 1x10^6 cells/sample were transferred into a 1.5ml Eppendorf tube. The samples were centrifuged at 200g for 8 mins at 4°C in a tabletop centrifuge. The supernatant was removed and 100μl ice-cold solvent mixture (methanol:acetonitrile:H2O (2:2:1, v/v) MS-grade; cooled to − 20 °C) was added to the pellet. The cells were mechanically homogenized with a P1000 pipett for 1 min followed by sonication for five minutes in a water bath sonicator at RT. Samples were further processed as described for the untargeted metabolomics analysis and stored at -80C until further processed. <u>Sample analysis for untargeted metabolomic profiling:</u> Extracts were thawed on ice, centrifuged for 5 min at 15,000 g, and directly injected onto the respective phase system. 10 μL of each sample were pooled and used as a quality control sample (QC). Samples were randomly assigned into the autosampler and injected on the respective phase system. For HILIC (hydrophilic interaction chromatography), an iHILIC®-(P) Classic, PEEK column, (100mm x 2.1mm, 5μm) with a precolumn (HILICON, Umeå, Sweden) was used. Here, a 26 min gradient from 90% A (acetonitrile) to 80% B (25 mM ammonium bicarbonate in water) was used, employing a flow rate of 100 μL/min delivered through an Ultimate 3000 HPLC system (Thermo Fisher Scientific, Germany). Following analysis by HILIC-MS/MS, samples were analyzed with reversed phase chromatography. Here, an ACQUITY UPLC HSS T3 column (150 mm x 2.1 mm; 1.8 μm) with VanGuard precolumn (Waters Corporation, Millford, MA, USA) was used. A 20 min gradient of 99% A (0.1% formic acid in water) to 60% B (acetonitrile with 0.1% formic acid) has been employed using the same HPLC system and flow rate. In both cases, metabolites were ionized via electrospray ionization in polarity switching mode after HILIC separation and in positive polarity mode after RP separation. Sample spectra were acquired by data-dependent high-resolution tandem mass spectrometry on a Q-Exactive Focus (Thermo Fisher Scientific, Germany). The ionization potential was set to +3.5/-3.0 kV, the sheet gas flow was set to 20, and an auxiliary gas flow of 5 was used. Samples were analyzed in a randomized fashion and QC samples were additionally measured in confirmation mode to obtain additional MS/MS spectra for identification. Obtained data sets have been processed by compound discoverer 3.0 (Thermo Fisher Scientific). Compound annotation was with a mass accuracy of 3 ppm for precursor masses and 10 ppm for fragment ion masses search public spectral databases as well as our in-house spectral library. Here, also experimentally obtained retention times have been used for the validation of metabolite identifications.

<u>Sample analysis for targeted metabolic profiling</u>: Polar metabolites have been analyzed using HILIC-LC-MS/MS. Each sample was injected onto an iHILIC®-(P) Classic, PEEK column, (100mm x 2.1mm, 5μm) with a precolumn (HILICON, Umeå, Sweden). An Ultimate 3000 HPLC system (Dionex, Thermo Fisher Scientific) has been used, employing a flow rate of 100 μl/min and directly coupled to a TSQ Quantiva mass spectrometer (Thermo Fisher Scientific). A 15-minute gradient from 14% B to 80% B (A: 95% acetonitrile 5% 10 mM aqueous ammonium acetate; B: 5 mM aqueous ammonium bicarbonate) has been used for separation. The following transitions have been used for quantitation in the negative ion mode (2.8 kV): pyruvate: 87 m/z → 43 m/z, lactate: 89 m/z → 43 m/z, taurine: 124 m/z → 80 m/z, ketoleucine: 129 m/z → 85 m/z, α-ketoglutaric acid: 145 m/z → 101 m/z, AMP: 346 m/z → 79 m/z, IMP: 347 m/z → 79 m/z, ADP: 426 m/z → 134 m/z, ATP: 506 m/z → 159 m/z, NAD: 662 m/z → 540 m/z, NADH: 664 m/z → 408 m/z, NADP: 742 m/z → 620 m/z, NADPH: 744 m/z → 426 m/z, CoA: 766 m/z → 408 m/z, Acetyl-CoA: 808 m/z → 408 m/z and in the positive ion mode (3.5kV) GSH: 308 m/z → 408 m/z, GSSG: 613 m/z → 355 m/z and SAM: 399 m/z → 250 m/z. The remaining metabolites have been quantified by reverse phase LC-MS/MS, injecting 1 ul of the metabolite extract onto a RSLC ultimate 3000 (Thermo Fisher Scientific) directly coupled to a TSQ Altis mass spectrometer (Thermo Fisher Scientific) via electrospray ionization. A Kinetex C18 column was used (100 Å, 150 x 2.1 mm), employing a flow rate of 80 μl/min. A 7-minute-long linear gradient was used from 99% A (1 % acetonitrile, 0.1 % formic acid in water) to 60% B (0.1 % formic acid in acetonitrile). Liquid chromatography-tandem mass spectrometry (LC-MS/MS) was performed by employing the selected reaction monitoring (SRM) mode of the instrument in the positive ion mode, using the transitions 156 m/z → 110 m/z (histidine), 175 m/z → 70 m/z (arginine), 241 m/z → 74 m/z (cystine), 76 m/z → 30 m/z (glycine), 133 m/z → 70 m/z (ornithine), 175 m/z → 74 m/z (asparagine), 106 m/z → 60 m/z (serine) 120 m/z → 74 m/z (threonine), 147 m/z → 84 m/z (lysine), 147 m/z → 130 m/z (glutamine), 148 m/z → 84 m/z (glutamic acid) 90 m/z → 4 m/z (alanine and sarcosine), 104 m/z → 84 m/z (GABA), 176 m/z → 159 m/z (citrulline), 116 m/z → 70 m/z (proline), 118 m/z → 72 m/z (valine), 150 m/z → 133 m/z (methionine), 132 m/z → 86 m/z (isoleucine and leucine), 182 m/z → 136 m/z (tyrosine), 166 m/z → 103 m/z (phenylalanine), 205 m/z → 188 m/z (tryptophane), 134 m/z → 74 m/z (aspartic acid) 177 m/z →160 m/z (serotonin) and 154 m/z → 137 m/z (dopamine).

For all transitions, the optimal collision energy was defined by analyzing pure metabolite standards. LC-MSMS chromatograms were interpreted using TraceFinder (Thermo Fisher Scientific). After LC-MS/MS analysis retention times were verified by standard addition of pure compounds to arbitrarily selected samples, validating experimental retention times with the respective pure substances.

### Analysis of the untargeted metabolomics dataset

Code can be found in github repository: https://github.com/menchelab/novarinocollab Combining hydrophilic interaction and reversed phase chromatography results and filtering: Measurement of metabolites was acquired by LC-MS using hydrophilic interaction liquid chromatography (HILIC) or reversed phase chromatography (RP). Both of which contain overlapping metabolites. Assuming quantifications in RP would align with quantifications in HILIC, we combined HILIC and RP by adding only those metabolites that are not present in HILIC. Subsequently, the combined data was filtered to retain only those metabolites that were mapped to a KEGG ID. Principal component analysis: Principal component analysis was conducted in Python using scikit-learn v1.0.2 (https://dl.acm.org/doi/10.5555/1953048.2078195). Metabolite timecourse dynamics analysis: Timecourse dynamics analysis was conducted on the combined and filtered metabolomics data. In brief, the mean abundance of the metabolites in WT and KO at each timepoint was computed and resulting averages were normalized to values between 0 and 1 such that the maximum abundance over time is 1 and the minimum abundance over time is 0. This was done for each metabolite in each genotype separately. Differential dynamics of metabolites between the genotypes were assessed by computing the Pearson correlation coefficient r between normalized trajectories in WT and KO where metabolites with r < 0.975 were considered to show differing dynamics. Grouping of metabolite trajectories in dynamics classes: Normalized trajectories in WT and KO were grouped using a gaussian mixture model (GMM) with six components (implemented in scikit-learn v1.0.2 (https://dl.acm.org/doi/10.5555/1953048.2078195)). The optimal number of components was assessed with the Bayesian information criterion. The resulting clusters were aligned to HILIC WT clusters and RP and HILIC results were combined as described above only retaining metabolites with a mapped KEGG ID. Ternary plot generation: Raw timepoint values for each metabolite in the combined and filtered data were averaged over replicates and normalized to sum to 1. These values were then plotted using the python-ternary library v1.0.8 (https://github.com/marcharper/python-ternary). Metabolic pathway enrichment: For classical pathway enrichment analysis, list of pathway annotations of classes ‘metabolism’ and ‘information processing’ was retrieved from the Kyoto Encyclopedia of Genes and Genomes (KEGG). For the custom cluster enrichment analysis metabolites were manually grouped and assigned unique IDs. Enrichment analysis was conducted by assessing the significance of overlap between a given set of metabolites and a pathway with Fisher’s exact test and subsequent multiple testing adjustment using the Benjamini-Hochberg procedure using an FDR of 0.05. Circos plot generation: Circos plots to visualize correlations between metabolite groups were generated as described previously (Yu et al., 2020). In brief, the Pearson correlation coefficient r over all replicates was computed for each pair of metabolites and each pair of metabolite groups. Subsequently, only correlations of |r| > 0.5 were used to compute widths of plot elements. Finally, circus plots were generated using the pyCircos v0.3.0 (https://github.com/ponnhide/pyCircos)

### Proteomic analysis

<u>Sample collection:</u> P5 *Slc7a5^fl^;Emx1-Cre* littermates (n=5/genotype) were decapitated and the cortex was dissected on ice. After removal of the hippocampus, the exact tissue weight was determined. The samples were snap frozen until used for further steps. <u>Sample preparation for mass spectrometry:</u> All samples were processed with the iST-NHS kit from PreOmics GmbH. A modified protocol for on-beads digest (sonication for 10*30s in a Bioruptor sonicator in presence of 50 mg Protein Extraction beads, C20000021, Diagenode) was used for tissue samples followed by a 3.5h digestion step. A TMT-10plex kit was used (lot VK306785). For lot VK306785, each 0.8 mg vial was split into 5 parts: 3 parts were used to label 1 of the 10 tissue samples; 12 unrelated samples, as well as 3 aliquots of an additional 13th sample (mixed sample for Internal Reference Scaling), was labeled with 1 remaining part. Samples were cleaned up and combined. The whole lysate TMT sample was also fractionated by offline High pH Reversed Phase fractionation into 48 fractions (A: de-ionized water + 10 mM NH4OH; B: 90% LC-grade Acetonitrile + 10 mM NH4OH; flow: 0.15 mL/min; 0-4 min: 1% B, 115 min: 25%, 140 min: 40%, 148 min: 75%, maintained for 12 min, followed by 45 min equilibration at 1% B). Whole lysate fractions were vacuum-dried overnight and sent for MS analysis. <u>LC/MS-MS analysis:</u> All samples were analyzed by LC-MS/MS on an Ultimate 3000 nano-HPLC (Dionex) coupled with a Q-Exactive HF (ThermoFisher Scientific). Chromatographic method: peptide samples were loaded onto either a GEN1 uPAC column (Pharmafluidics, first 2 projects) or a 50 cm EasySpray column (ES903: 75 μm ID, 2 μm particle size, ThermoScientific); solvent A: H2O, 0.1% formic acid; solvent B: 80% acetonitrile in H2O, 0.08% formic acid; gradients: tissue fractions, 2% to 44% B in 60 min; Mass Spectrometry method: Data-Dependent acquisition (Full MS / dd-MS2); MS1: 1 microscan, 120,000 resolving power, 3e6 AGC target, 50 ms maximum IT, 380 to 1,500 m/z, profile mode; up to 20 data-dependent MS2 scans per duty cycle, excluding charges 1 or 8 and higher, dynamic exclusion window 10s (60 min gradient) or 60s (180 min gradients): isolation window 0.7 m/z, fixed first mass 100 m/z, resolving power 60,000, AGC target 1e5, maximum IT 100 ms, (N)CE 32. <u>Data analysis:</u> The dataset was searched in MaxQuant (1.6.17.0) against a Mus musculus fasta database downloaded from UniProtKB. Fixed modification was set to Carbamidomethyl (label-free samples) or C6H11NO (TMT samples). Variable modifications were set to include Acetyl (protein N-term), Oxidation (M), Gln->pyroGlu and Deamidation (NQ) and Phospho (STY). Match-between-runs and second peptides were set to active. All FDRs were set to 1% (tissue samples). Each MaxQuant output evidence.txt files was then re-processed separately in R using in-house scripts. Evidence reporter intensities were corrected using the relevant TMT lot’s purity table, scaled to parent peptide MS1 intensity and then normalized using the Levenberg-Marquardt procedure. The long format evidence table was consolidated into a wide format peptidoforms table, adding up individual values where necessary. Peptidoform intensity values were log10 transformed. Values were re-normalized (Levenberg-Marquardt procedure). Protein groups were inferred from observed peptidoforms, and, for each group, the estimated expression values across samples were calculated by averaging individual peptidoform log10 intensity vectors, scaling the vector to reflect the intensity level of the most intense peptidoform according to the best flyer hypothesis (phospho-peptides and their unmodified counterpart peptide were excluded). Peptidoform and protein group log2 ratios were calculated: to the average reference (WT) sample (whole lysate dataset). Statistical significance was tested with the limma package, performing both a moderated t-test and an F-test (limma package). The Benjamini-Hochberg procedure was applied to compute significance thresholds at various pre-agreed FDR levels. Regardless of test, protein groups with a significant p-value were deemed to be regulated if their absolute log2 fold change (= logFC) was greater than 95% of control to average control logFC. During the analysis phase two out of five mutant samples behaved as outliers and were excluded from further analysis. Protein groups were annotated with GO terms, applying a term if it, or one of its offspring terms, was found among the annotations of any protein accession which could explain all peptides in the group. GO enrichment analysis was performed using an in-house script built around the topGO package, comparing separately for each contrast all up- or down-regulated proteins, or both, against the background of all identified protein groups.

### Bulk RNA-Sequencing of *Slc7a5^fl^;Emx1-Cre* cortical tissue

<u>Sample preparation</u>: P1-P2 pups were decapitated and the brains were quickly dissected on ice under RNAse free conditions. Total RNA of one cortical hemisphere was extracted by using 500 μL Trizol (Thermo Fisher Scientific) for homogenization and 500 μL chloroform (Sigma-Aldrich), followed by centrifugation at 12000 g for 15 min at 4°C. The upper phase was transferred to a fresh tube and 1.5 volumes of 100% ethanol were added. Total RNA was purified by using the RNA Clean&Concentrator-5 prep Kit (Zymo Research). The samples were further treated with RQ1 RNase-Free DNase (Promega) as described in the kit instructions manual. RNA sample quality was assessed by using the NanoDrop spectrophotometer (Thermo Fisher Scientific) and the Bioanalyzer 2100 with the RNA 6000 Nano kit (Agilent). cDNA libraries were generated with the SENSE mRNA-Seq Library Prep Kit V2 (Lexogen) using 1.5mg total RNA. The quality of the generated libraries was monitored by using the High Sensitivity DNA Analysis Kit (Agilent) and the Bioanalyzer 2100. Libraries were sequenced on an Illumina HiSeq 2500 instrument. <u>Analysis</u>: Demultiplexed raw reads were trimmed before alignment using the FASTX toolkit. Trimmed reads were aligned to the mouse genome using STAR version 2.5.4 (genome: GrCm38, gene annotation: Gencode release M8). Read counts per gene were quantified using STAR. The aligned sequencing data were uploaded to the Galaxy web platform. Differential expression analysis was performed in usegalaxy.org using the Bioconductor package DESeq2 (PMID: 25516281) using an FDR threshold of 0.05. Gene Ontology enrichment analysis was performed using the Bioconductor package GOstats version 2.36.0 with a p value cutoff of 0.001 and conditional testing enabled. For bulk RNA sequencing n = 3 mice per genotype were employed.

### Electrophysiology

<u>Sample preparation</u>: Acute brain slices were obtained from P6-7 *Slc7a5^fl/fl^;Emx1-Cre* and *M8-TG-Sc7a5^fl/fl^/GT;Emx1-Cre*+ animals and wild type littermates of both sexes. Further, *Slc7a5^fl^;Emx1-Cre* animals were also used at P25-P26. Coronal sections (300 μm) were prepared from primary somatosensory cortex. Animals were decapitated under isoflurane anesthesia and whole brains were rapidly removed from the skull and sectioned using a VT 1200S vibratome (Leica Microsystems) in ice-cold cutting solution, containing (mM): 87 NaCl, 25 NaHCO3, 2.5 KCl, 1.25 NaH2PO4, 10 glucose, 75 sucrose, 7 MgCl2, 0.5 CaCl2 (320 mOsm, 7.2-7.4 pH). When older mice were tested (P25-26), slices were sectioned in ice-cold cutting solution, containing (mM): 93 NMDG, 2.5 KCl, 1.2 NaH2PO4, 30 NaHCO3, 20 HEPES, 25 glucose, 5 sodium ascorbate, 2 thiourea, 3 sodium pyruvate, 10 MgCl2, 0.5 CaCl2 (320 mOsm, 7.2-7.4 pH). Slices from P25-26 mice were recovered at 32°C for 10-12 minutes in the same solution and then allowed to recover at room temperature for at least 1 hour in regular artificial cerebrospinal fluid (ACSF), containing (mM): 125 NaCl, 2.5 KCl, 1.25 NaH2PO4, 25 NaHCO3, 25 glucose, 1 MgCl2 and 2 CaCl2 (320 mOsm, 7.2–7.4 pH). The ACSF was continuously oxygenated with 95% O2 and 5% CO2 to maintain the physiological pH. Slices were visualized under infrared-differential interference contrast (IR-DIC) using a BX-51WI microscope (Olympus) with a QIClickTM charge-coupled device camera (Q Imaging Inc, Surrey, BC, Canada). <u>Recordings</u>: Patch pipettes (4-6 MΩ; World Precision Instruments) were pulled on a P-1000 puller (Sutter Instruments) and filled with the intracellular recording solution, containing (mM): 128 K-gluconate, 10 HEPES, 10 Na2-phosphocreatine, 1.1 EGTA, 5 MgATP, 0.4 NaGTP (osmolarity adjusted to 295 mOsm with sucrose, 7.3-7.4 pH). Current clamp recordings were performed at room temperature (24 ± 1 °C) from layer II/III pyramidal neurons. When experiments were performed in *M8-TG-Sc7a5/GT;Emx1-Cre*+ animals, patch clamp from layer II/III was visually guided by fluorescent labeling of neurons to recognize wild type (red) and knock-out (green) pyramidal neurons in the same brain slice. <u>Analysis</u>: Membrane capacitance and resting membrane potential were determined immediately after the establishment of whole-cell configuration. Neuronal membrane potential was held at approximately -60 mV (P6-7 mice) or -70 mV (P25-26 mice) by constant current injection. Current steps ranging in amplitude from -40 to +50 pA (10 pA increments; 600 ms duration) were applied to estimate the *f – I* relationship. In current clamp experiments from P25-26 mice, current steps ranging in amplitude from -300 to +400 pA (50 pA increments; 600 ms duration) were applied to estimate the *f – I* relationship. Properties of individual action potentials (APs) were determined from the first current step necessary to elicit at least one AP. Phase-plane plot analysis was performed to evaluate the dynamic changes of the membrane potential over time (dV/dt). The threshold was set as the voltage at which the first derivative of the voltage trace reached 20 V/s. Amplitude was calculated as the difference between the threshold and the peak. AP half-width was measured at half the difference between the firing threshold and the AP peak. Current clamp recordings were filtered at 2 kHz, sampled at 20 kHz and acquired using a MultiClamp 700B amplifier and a Digidata 1550A. Recorded signals were analyzed off-line using the Clampfit 10 software (Molecular Devices).

### Behavioral analysis

All behavioral tests were performed during the light period. Mice were habituated to the test room 24hr before each test. For all studies sex-matched littermates were employed. Equipment was cleaned between each trial with 70% ethanol. In order to recover, mice were given 24hr between different tests. All behavioral studies were performed starting with the least aversive task first and ending with the most aversive one. Behavioral tests were carried out with P55 to P65 animals. <u>Open field test</u>: Exploratory behavior in a novel environment was assessed in an open field arena (45cmL x 45cmW x30cmH) made out of dark Plexiglas. The animal was placed in the center of the arena and recorded for 20 min. Locomotor activity (distance traveled and velocity) in the center/periphery of the arena as well as rearing were tracked and analyzed using the EthoVision XT 11.5 software (Noldus). <u>Three chamber sociability test</u>: Mice were tested for social deficits as described previously (Moy et al., 2007). In detail, the behavior of the animals was monitored in a rectangular three chambers arena (60cm (L) x 40cm (W) x 20 cm (H)) made of clear Plexiglas. Age and sex matched littermates were used for all tests. Sex- and age-matched C57BL/6J mice were used as “stranger” mice. They were habituated to the wire cage for 2x 10 min 24hr prior to the test. During the first session (habituation) each subject was placed into the center chamber with open access to both left and right chamber, each chamber containing an empty wire cage. 10 min of habituation were followed by the social phase. Thereby, an age-matched stranger was placed in the wire cage of the left chamber while a novel object was placed into the right chamber’s cage. The wire cage (12cmH, 11cm diameter) allows nose contact between the test subject and the C57BL/6J strangers. The test animal was allowed to freely explore the arena for 10 min. Locomotor activity (distance traveled and velocity) the number of nose contacts (< 5cm proximity) with the caged mouse/object was recorded and analyzed by EthoVision XT 11.5 software (Noldus). Vertical explorative behavior was assessed by manually quantifying the number of rearings during the habituation phase. <u>Gait measurement test</u>: Potential gait impairments were monitored by using “the footprint test”, as previously described (Carter et al., 2001). In detail, fore and hind paws were painted with dyes of contrasting colors. The mouse was placed in a narrow corridor on white paper. A darkened house was used as a bait to encourage the mouse to walk in a straight line. The footprint patterns were then analyzed for stride length, sway length and stance length. <u>Hind limb clasping test</u>: To assess potential hind limb clasping behavior, mice were suspended by their tails for 10s. During this period the hind limb position was monitored and scored according to the severity of the phenotype (Guyenet et al., 2010). The test was repeated three times for each animal.

## ACKNOWLEDGEMENTS

We thank A. Freeman, V. Voronin, and E. Novak for technical assistance, S. Deixler and A. Stichelberger for the management of our animal colony, as well as M. Schunn and the Preclinical Facility team for technical assistance. We thank L. Andersen and J. Sonntag, who were involved in the generation of the MADM lines. We thank A. Nicolas and the ISTA proteomic facility for technical assistance in the proteomic analysis, as well as the ISTA electron microscopy and ISTA Imaging and Optics facility for assistance and technical support. Metabolomics LC-MS/MS analysis was performed by the Metabolomics Facility at Vienna BioCenter Core Facilities (VBCF), member of the Vienna BioCenter (VBC), Austria and funded by the City of Vienna through the Vienna Business Agency. Bulk RNA Sequencing was performed by the Next Generation Sequencing Facility at Vienna BioCenter Core Facilities (VCBF). Schematics were generated using Biorender.com. This work was supported by the Austrian Science Fund (FWF, DK W1232-B24) and by the European Union’s Horizon 2020 research and innovation program (ERC) grant 725780 (LinPro) to S.H. and 715508 (REVERSEAUTISM) to G.N..

## AUTHOR CONTRIBUTIONS

L.S.K., B.B., L.A.S., S.G. and N.A. performed experiments. D.M. performed the analysis of the metabolomics and proteomics datasets. G.N. conceived and supervised the study. M.G.B. and M.S. recruited and characterized patients. N.A., F.P., T.R. and S.H. were involved in the generation of the MADM lines. G.N. wrote the paper together with L.S.K.. All authors read and approved the final version of the manuscript.

---

**Figure S1.**
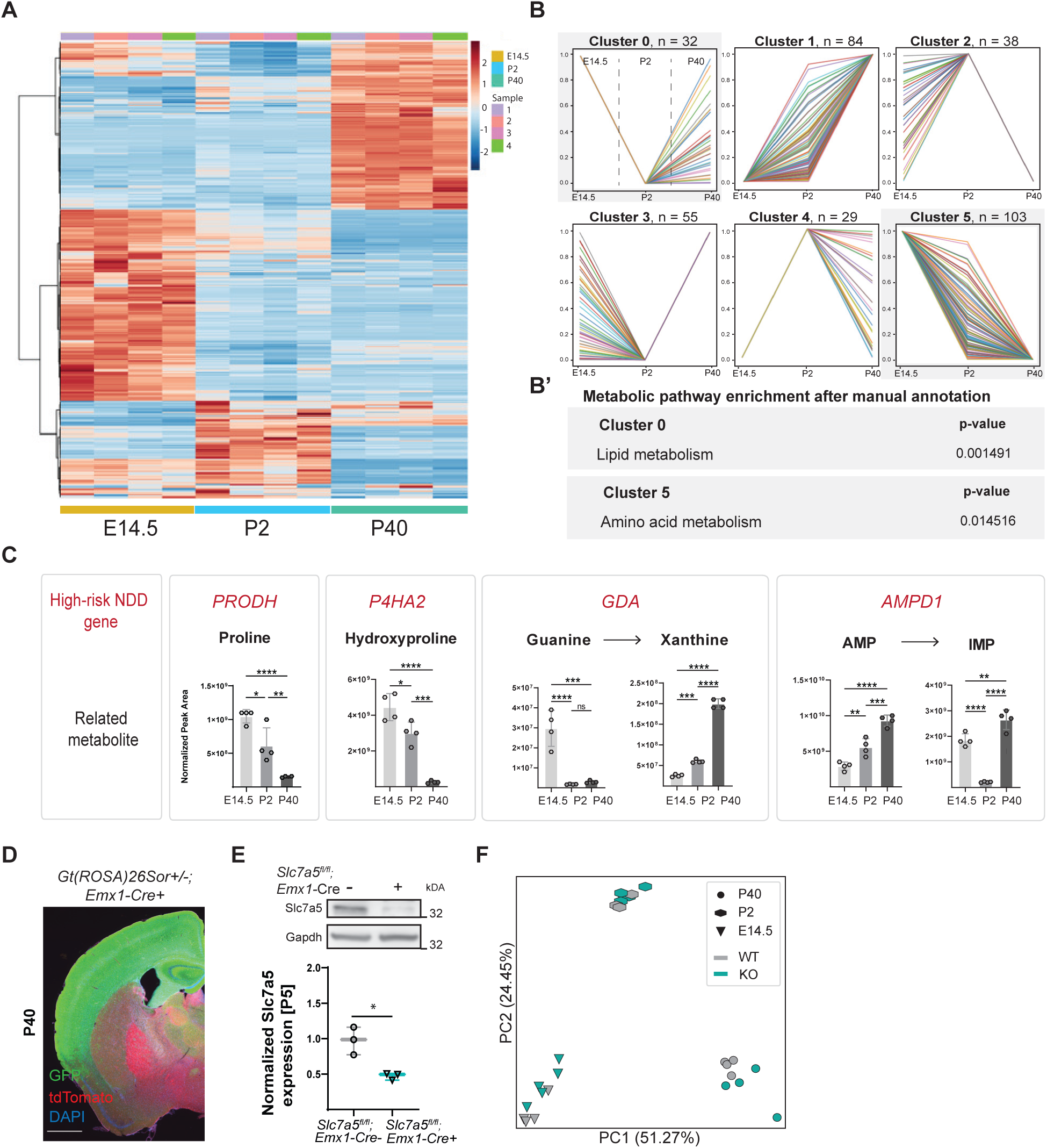
Untargeted metabolomic profiling of wild-type and *Slc7a5* deficient cortex. (**A**) Heatmap visualizing changes in metabolite levels in the wild-type cortex obtained from multiple mice at three developmental time points (E14.5, P2, P40). (**B)** Clustering of all the detected metabolites based on their developmental trajectory (**B**’) Metabolic pathway enrichment analysis of the different clusters employing literature-curated annotations (Amino acid, lipid and nucleoside metabolism, bioenergetic processes, glycolysis, others; Supplementary data 7) (**C**) Examples of metabolism-associated genes linked to neurodevelopmental disorders (**top**) and the relative levels of the associated metabolite(s) in the mouse cortex across time (**bottom**) (means ± SEM; n=4 animals per genotype and time point;*\*\*\*\*p < 0.00001*, *\*\*\*p < 0.0001*, *\*\*p < 0.01*; *\*p < 0.05; ^ns^p > 0.05*; 1-way ANOVA). (**D**) *Emx1*-driven Cre recombinase expression in neural cells of the neocortex was verified by utilizing the Gt26Sor^mtmG^ reporter mouse line. Gt26Sor^mtmGrep^;-Cre+ mice express tdTomato in all cells prior to Cre recombinase exposure. After recombination, Cre recombinase expressing cells are labeled with cell membrane-localized green fluorescent protein (GFP) (scale bar: 1500μm). (**E**) Western blot analysis of Gapdh normalized Slc7a5 expression levels of mutant and control cortical lysates (n=3 animals per genotype; *\*p < 0.05,* unpaired two-tailed *t*-test). (**F**) Principal component analysis of untargeted metabolomics analysis of *Slc7a5* mutant and wild-type cortex of all three time points.

**Figure S2.**
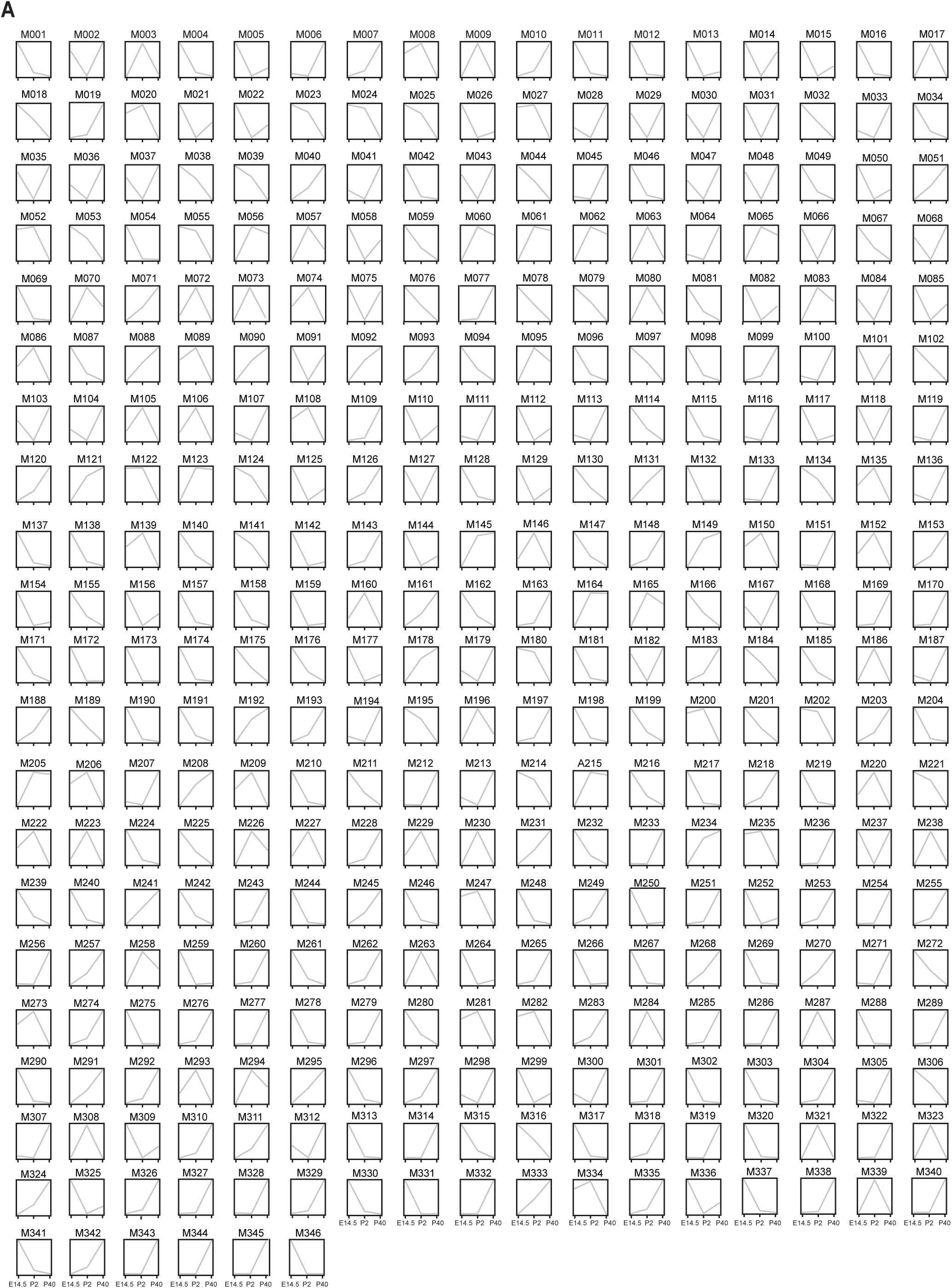
Developmental trajectories of metabolites detected in wild-type cortex. (**A**) Normalized and scaled trajectories of all metabolites detected in wild-type cortical tissue over time (x-axis: age (E14.5, P2, P40); y-axis: scaled abundance; metabolites: Supplementary data 1).

**Figure S3.**
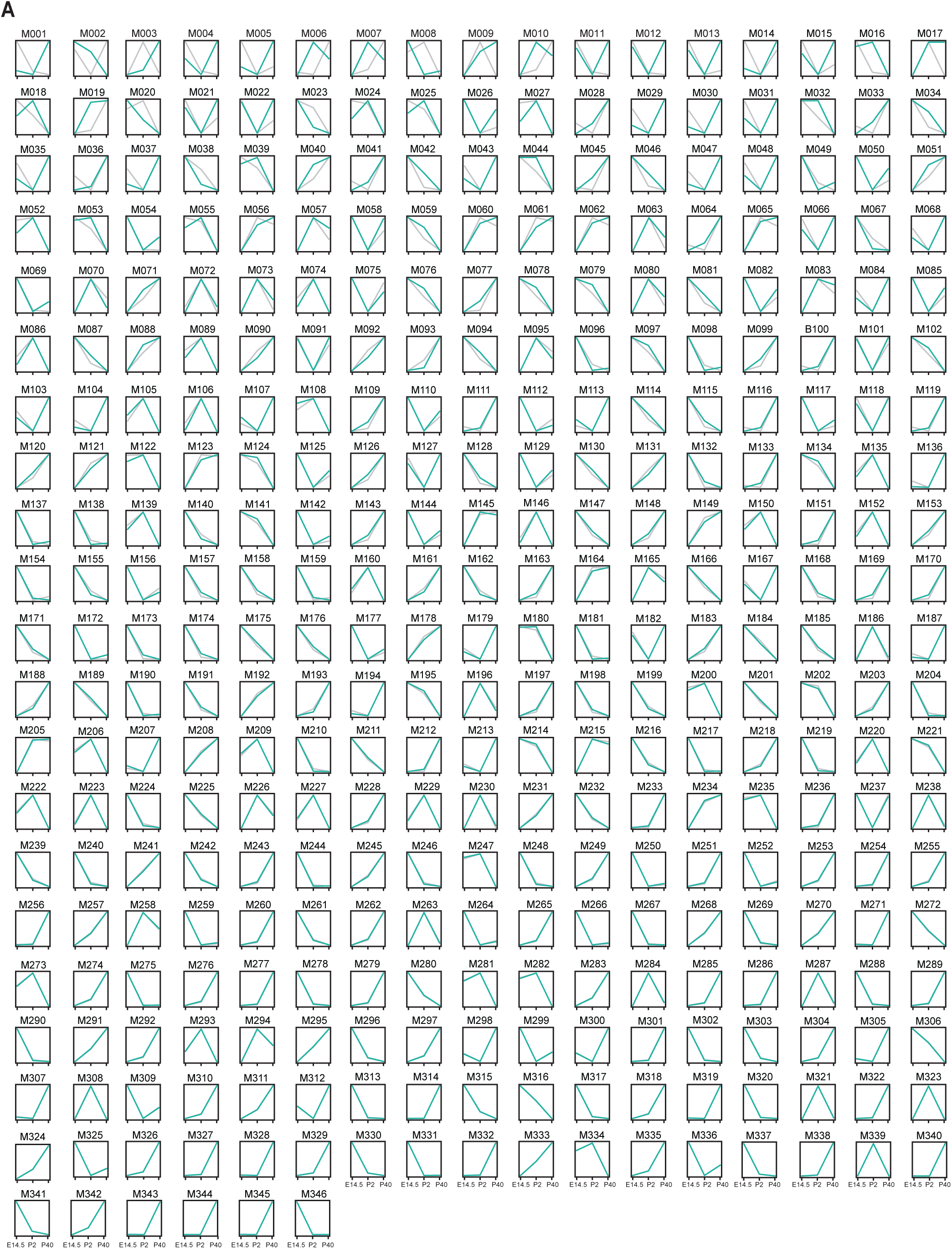
Developmental trajectories of metabolites detected in *Slc7a5* deficient cortex. (**A**) Normalized and scaled trajectories of all metabolites detected in *Slc7a5^fl/fl^;Emx1-Cre+* (cyan) and wild-type (gray) cortical tissue (x-axis: age (E14.5, P2, P40); y-axis: scaled abundance; metabolites and Pearson’s Coefficient: Supplementary data 1).

**Figure S4.**
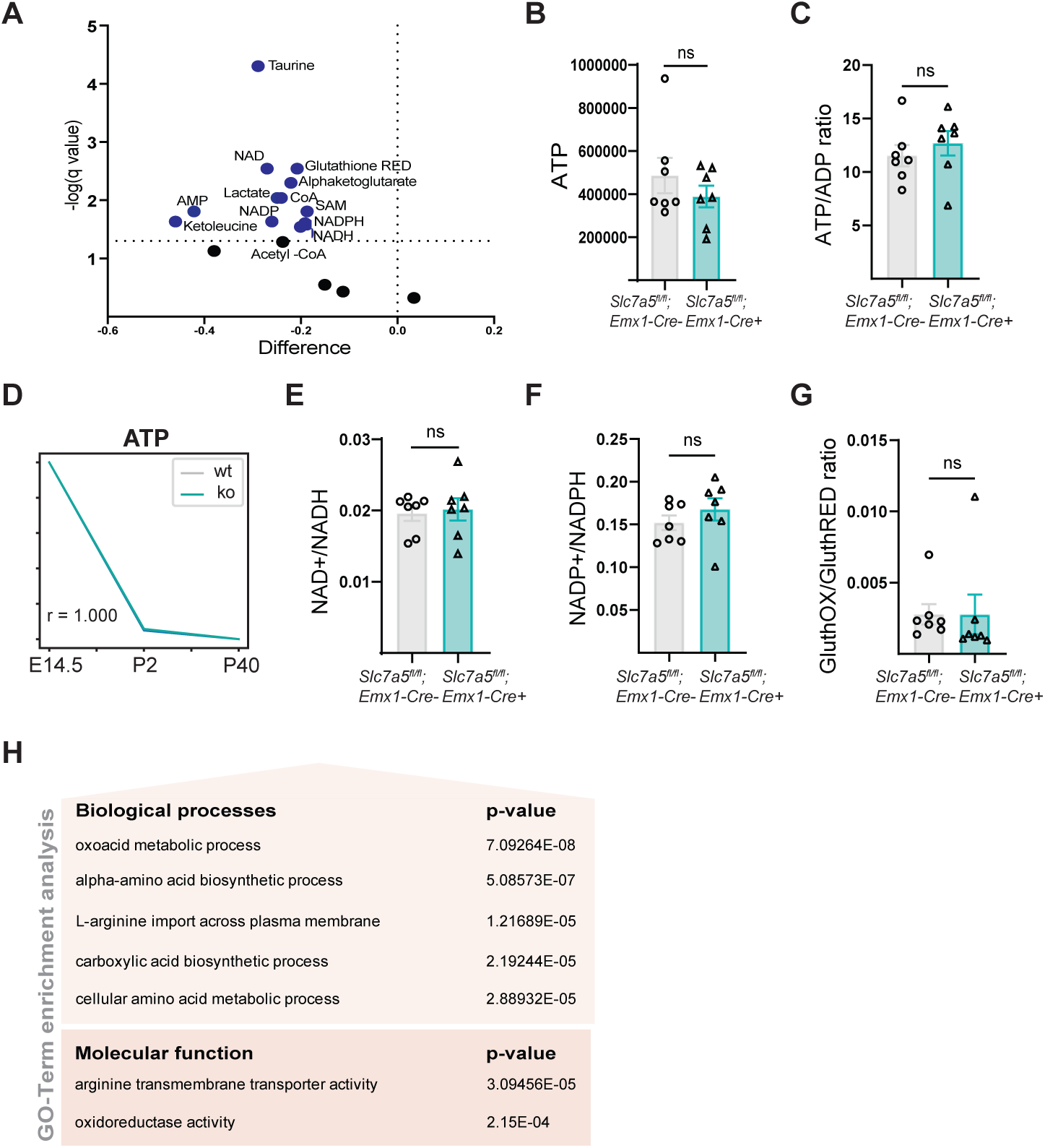
Cellular energy homeostasis is unaffected by the loss of *Slc7a5.* (**A**) Volcano plot of intracellular levels of metabolic co-factors and key metabolites of ATP producing pathways (n=7 mice per genotype; *^ns^p>0.05;* unpaired two-tailed *t-*test). Quantification of intracellular (**B**) ATP (means ± SEM; n=7 mice per genotype; FDR cut-off:1.0%), (**C**) ATP/ADP ratio (means ± SEM; n=7 mice per genotype; *^ns^p>0.05;* unpaired two-tailed *t-*test) and (**D**) ATP levels over the course of development in cortical tissue of *Slc7a5^fl/fl^;Emx1-Cre+* and wild-type mice at P2 (n=5 mice per genotype per time point; Pearson’s coefficient: r=1). (**E**) Intracellular NAD+/NADH and (**F**) NADP+/NADPH ratios are not changed in cortical cells of mutant and control mice (means ± SEM; n=7 mice per genotype; *^ns^p>0.05;* unpaired two-tailed *t-*test). (**G**) The ratio of oxidized and reduced Glutathione is unaffected in cortical cells of mutant mice compared to wild-type littermates (means ± SEM; n=7 mice per genotype; *^ns^p>0.05;* unpaired two-tailed *t-*test). (**H**) GO-term enrichment analysis of up-regulated genes of bulk RNA sequencing of *Slc7a5^fl/fl^;Emx1-Cre+* cortex at P2 (Selected GO-terms: Supplementary data 12; n=5 mice per genotype).

**Figure S5.**
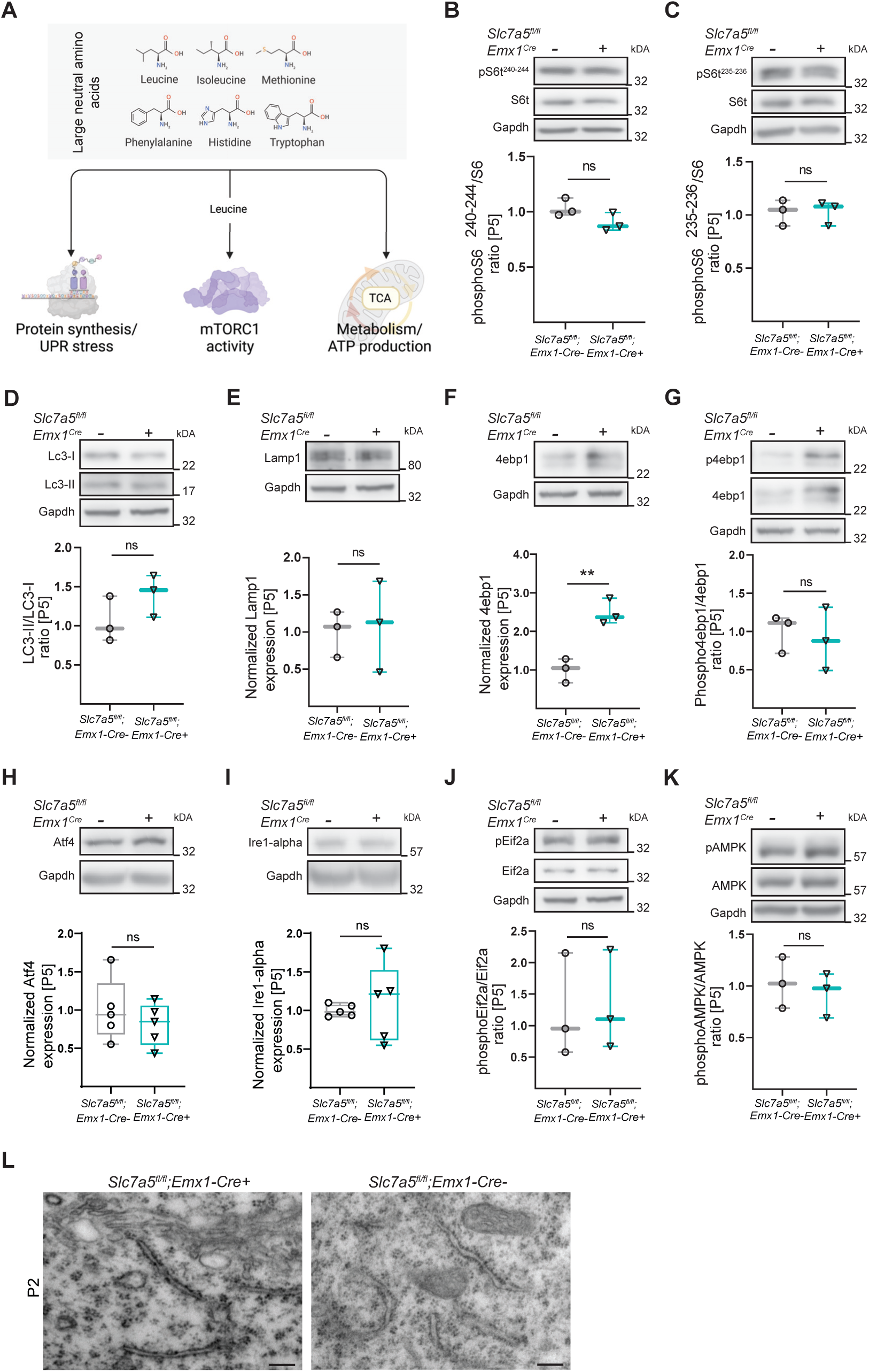
The mTOR, UPR and AMPK pathways are not affected in *Slc7a5* deficient mice. (**A**) Scheme of the signaling pathways and cellular processes which are linked thought to be link to large neutral amino acid levels. (**B-G**) Western blot analysis of markers used to quantify the state of the mTOR pathway and autophagy in P5 *Slc7a5^fl/fl^;Emx1-Cre-* and *Slc7a5^fl/fl^;Emx1-Cre+* cortices. Ratio of normalized (**B**) phosphoS6^240-244^/S6, **(C)** phosphoS6^235-236^/S6, (**D**) LC3I/II protein levels and (**E**) normalized Lamp1. Normalized (**F**) 4ebp1 expression and (**G**) phospho4ebp1/4ebp1 ratio (n = 3 mice per genotype; *^ns^p>0.05; **p<0.01;* unpaired two-tailed *t-*test). (**H-J**) Western blot analysis of markers used to monitor the unfolded protein response (UPR) pathway. Quantification of **(H**) Atf4, (**I**) Ire-alpha expression levels and (**J**) phosphoEif2a/Eif2a ratio normalized to Gapdh (n=4 mice per genotype; *^ns^p>0.05*). (**K**) Normalized phosphoAMPK/AMPK ratio (n=3 mice per genotype; *^ns^p>0.05;* unpaired two-tailed *t-*test). (**L**) Electron microscopy images of the endoplasmic reticulum (ER) showing unchanged ER morphology in mutant LII/III pyramidal neurons of P2 mice (scale bar: 24000x).

**Figure S6.**
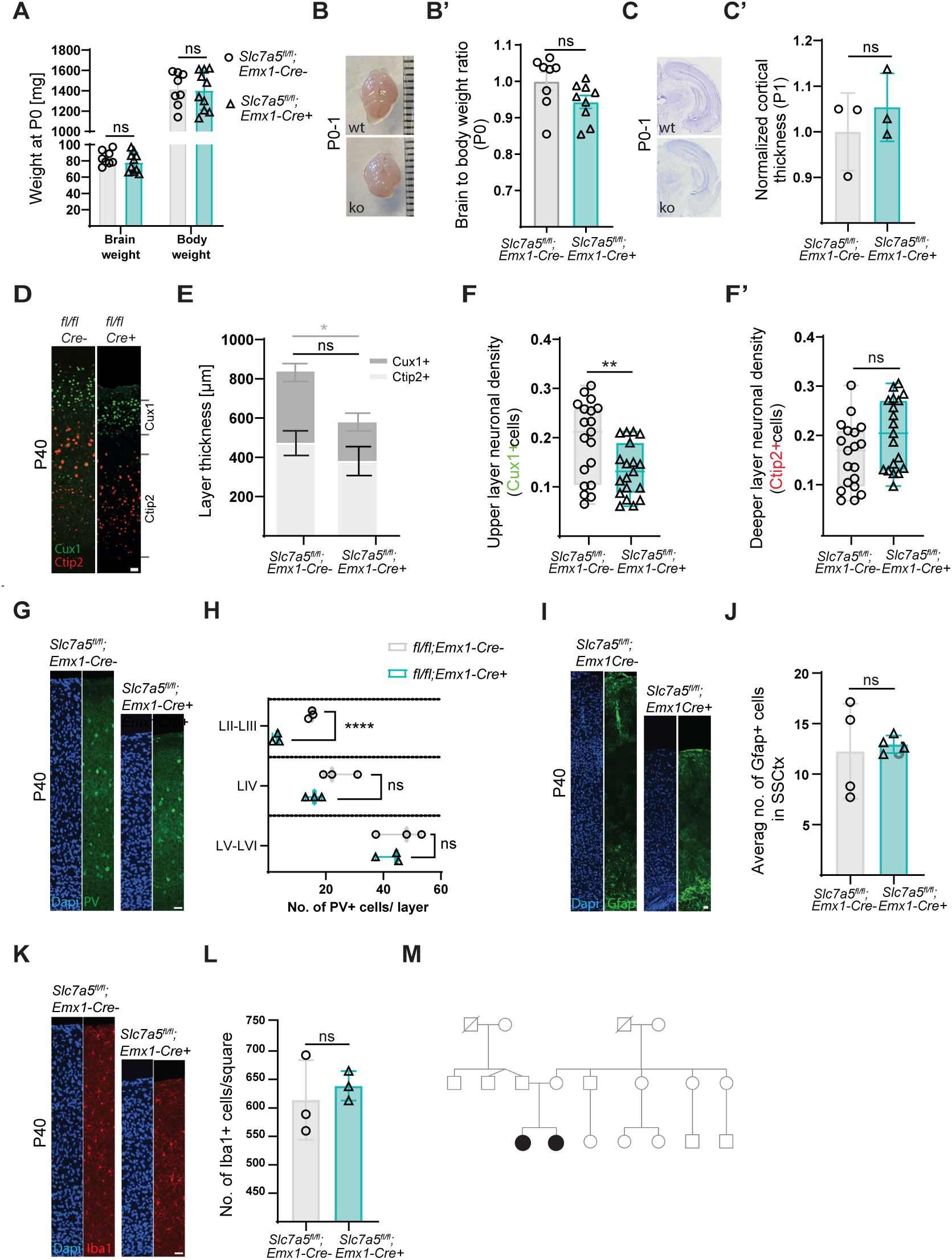
Characterization of the morphology and the cell-type composition of *Slc7a5*-deficient cortex. (**A**) Brain and body weight of *Slc7a5^fl/fl^;Emx1-Cre+* and wild-type littermates after birth (means ± SEM; n=8 animals per genotype; *^ns^p > 0.05,* unpaired two-tailed *t*-test). (**B**) Images and (**B’**) quantification of brain to body weight ratio of newborn (P0-P1) *Slc7a5^fl/fl^;Emx1-Cre+* and wild-type mice (means ± SEM; n=9 animals per genotype; *^ns^p > 0.05*; unpaired two-tailed *t*-test; scale: 1mm). (**C**) Nissl staining and (**C’**) quantification of cortical thickness in newborn mutant and wild-type mice (means ± SEM; n=3 animals per genotype; *^ns^p > 0.05*; unpaired two-tailed *t*-test). (**D**) Immunostaining for upper (Cux1) and lower (Ctip2) cortical layers in adult *Slc7a5^fl/fl^;Emx1-Cre* mice (scale bar: 100μm). (**E-F’**) Quantification of layer thickness (**E**) and cell density (**F-F’**) in Cux1+ or Ctip2+ cell layers (means ± SEM; n= 3 animal per genotype; n= 19 quantification squares for the cell density; *\*\*p < 0.01*; *^ns^p > 0.05;* unpaired two-tailed *t*-test). (**G**) Immunostaining and (**H**) quantification of the number of inhibitory (parvalbumin+) neurons in the different layers of cortical columns of adult mutant and wild-type mice (scale bar: 100μm; n=3 animals per genotype; *\*\*\*\*p < 0.0001*; *^ns^p < 0.05;* unpaired two-tailed *t*-test). (**I**) Immunostaining and (**J**) quantification of astrocytes (Gfap+) in cortical columns of adult mutant and wild-type mice (scale bar: 100μm; means ± SEM; n=4 animals per genotype; *^ns^p > 0.05*; unpaired two-tailed *t*-test; (**K**) Immunostaining and (**L**) quantification of microglia (Iba1+) cells in cortical columns of adult mutant and wild-type mice (scale bar: 100μm; means ± SEM; n=3 animals per genotype; *^ns^p > 0.05*; unpaired two-tailed *t*-test). (**M**) Pedigree displays a non-consanguineous background; two affected patients (solid symbols), and unaffected members (open symbols).

**Figure S7.**
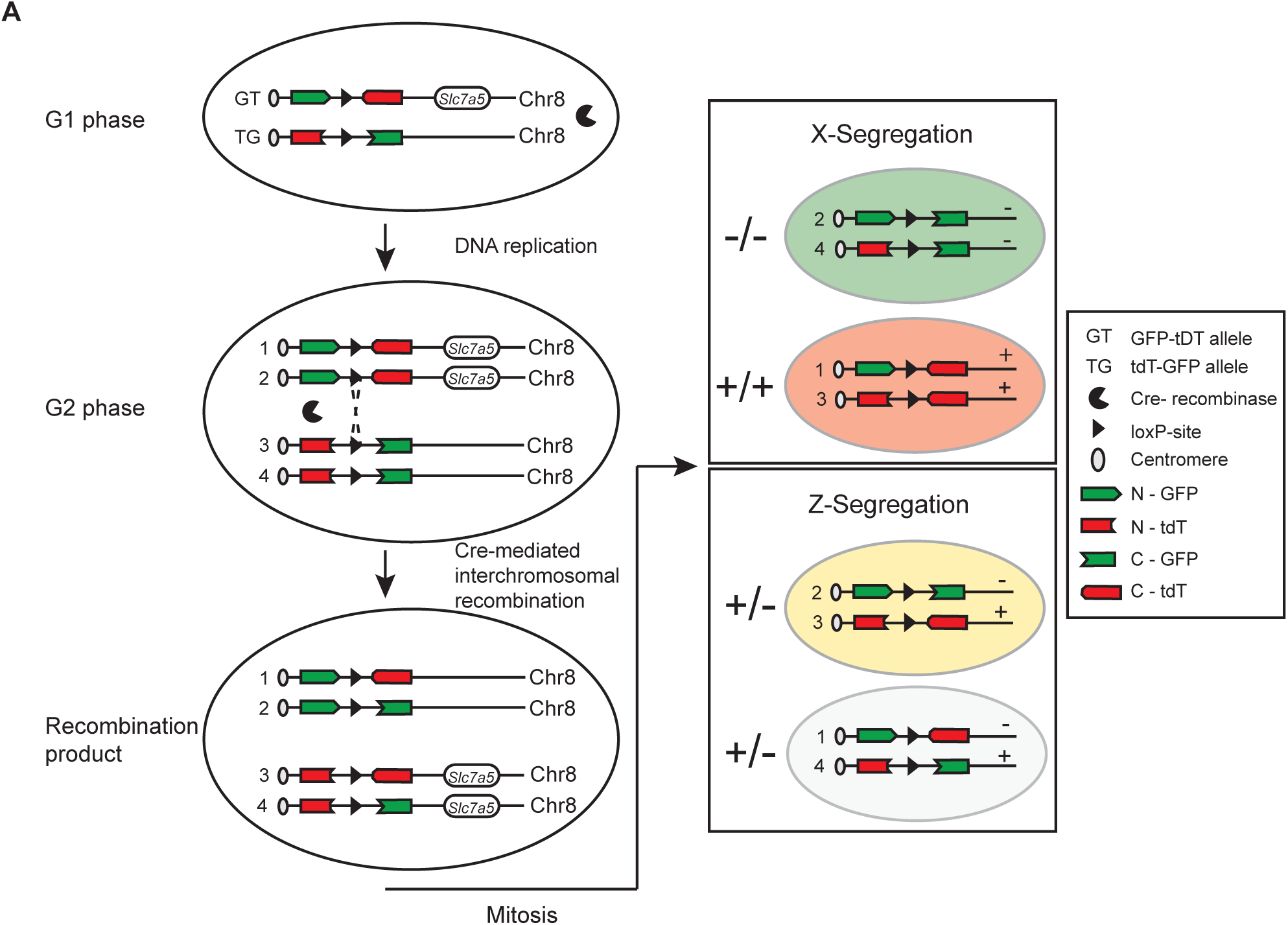
The mosaic analysis with double markers (MADM) principle. (**A**) Schematic of the MADM technique. Two reciprocally chimeric marker genes (MADM-8-cassettes) are inserted at two identical loci close to the centromeres distal to the *Slc7a5* gene on chromosome 8. Each cassette consists of two split coding sequences of green fluorescent protein (eGFP) and red fluorescent protein (tdTomato; cassettes are referred to as GT and TG). The N- and C-terminals of each reporter gene are separated by an intron containing a loxP site. This ensures that the chimeric genes do not produce functional proteins in the absence of Cre recombinase. In the presence of Cre recombinase, cis-recombination induces the deletion of the *floxed* exon in the *Slc7a5* gene, thereby generating a *Slc7a5*-knock out. These recombination events can take place throughout all phases of the cell cycle. In G2, recombination in trans can mediate stochastic interchromosomal recombination events at the loxP sites of the MADM cassettes. This restores functional eGFP and tdTomato expression in sparse single cells. During mitosis, two potential types of chromosomal segregation can take place. X-segregation generates green daughter cells homozygous for the mutation (Slc7a5^-/-^) and red homozygous for the wild-type allele (Slc7a5^+/+^), thereby creating fluorescently labeled genetic mosaic mice. Z-segregation produces one daughter cell that resembles the parental cell (colorless) and a second daughter cell expressing both fluorescent proteins (double colored). Both cells are heterozygous for *Slc7a5* mutation.

**Figure S8.**
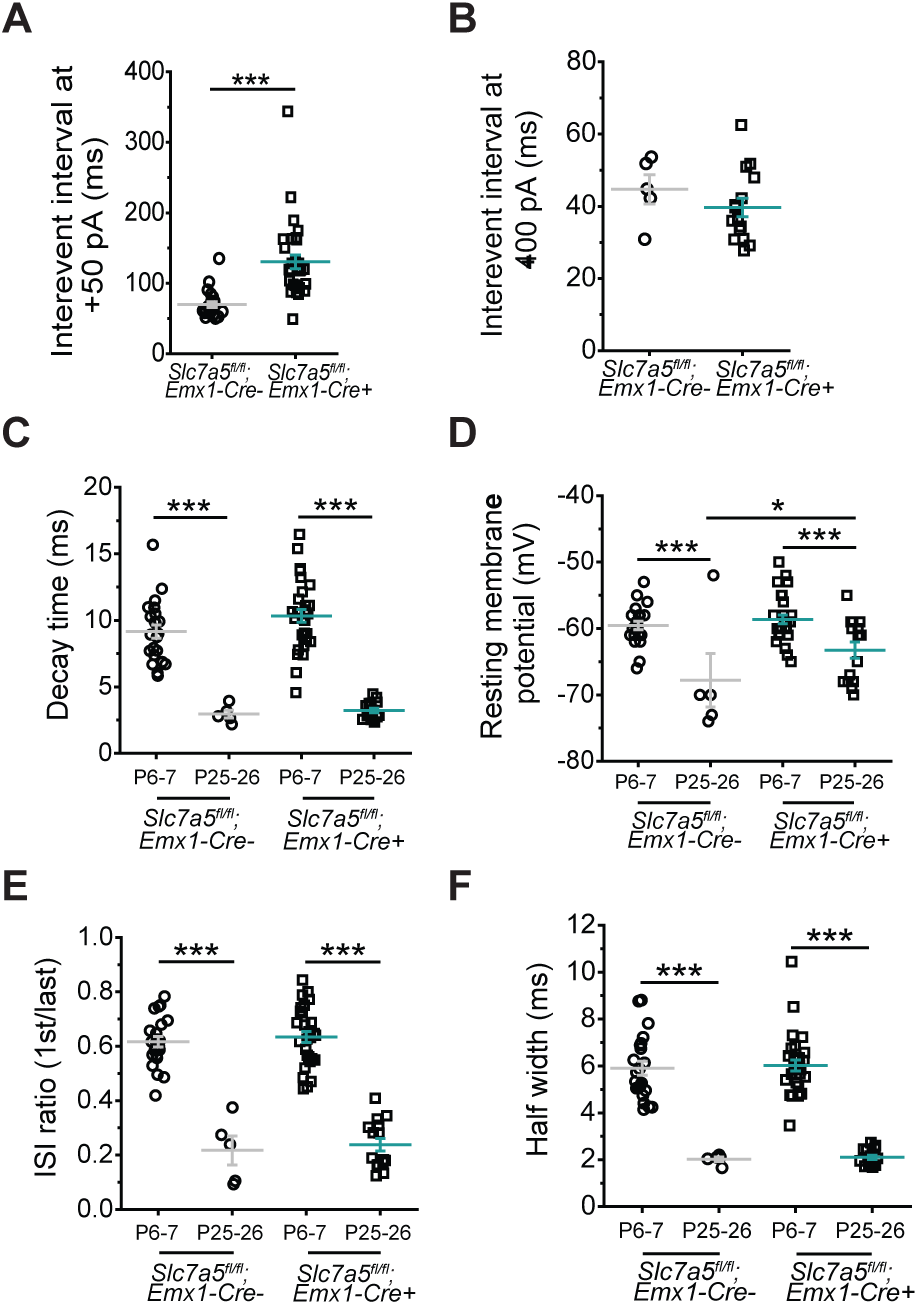
Electrophysiological properties of Slc7a5^fl/fl^;Emx1-Cre mice. (**A-B**) Inter-spike interval measured in current clamp experiments from P6-P7 (**A**) and P25-P26 (**B**) LII/III pyramidal neurons of *Slc7a5^fl/fl^;Emx1-Cre+* and *Emx1-Cre-* animals (*Emx1-Cre-*: n = 22 cells / 3 mice; *Emx1-Cre+*: n = 30 cells / 3 mice (P6-7); *Emx1-Cre+*: n = 5 cells / 3 mice; *Emx1-Cre-*: n = 15 cells / 3 mice (P25-26); ***p<0.001, unpaired two-tailed *t*-test). (**C**) Decay time, (**D**) resting membrane potential, (**E**) ISI ratio and (**F**) half width are not affected (*Emx1-Cre-*: n = 22 cells / 3 mice; *Emx1-Cre+*: n = 30 cells / 3 mice (P6-7); *Emx1-Cre-*: n = 5 cells / 3 mice; *Emx1-Cre+*: n = 15 cells / 3 mice (P25-26). Two-way ANOVA for AP decay time: genotype F(1,71) = 1.28 ^ns^p>0.5, time point F(1,71) = 107.51 ***p<0.001, interaction F(1,71) = 0.5 ^ns^p>0.5, Holm-Sidak post hoc ***p<0.001. Two-way ANOVA for resting membrane potential: genotype F(1,70) = 4.89 *p<0.5, time point F(1,70) = 20.35 ***p<0.001, interaction F(1,70) = 2.14 ^ns^p>0.5, Holm-Sidak post hoc *p<0.05 ***p<0.001. Two-way ANOVA for ISI ratio: genotype F(1,70) = 0.42 ^ns^p>0.5, time point F(1,70) = 181.02 ***p<0.001, interaction F(1,70) = 0.002 ^ns^p>0.5, Holm-Sidak post hoc ***p<0.001. Two-way ANOVA for half width: genotype F(1,71) = 0.08 ^ns^p>0.5, time point F(1,71) = 132.15 ***p<0.001, interaction F(1,71) = 0.002 ^ns^p>0.5, Holm-Sidak post hoc ***p<0.001.

**Table S1.** Overview of previously published and novel patients with biallelic pathogenic variants in the *SLC7A5* gene.

| Patient-ID | 1426-5 | 1426-6 | 1426-8 | 1426-19 | 1465-3 | 1465-4 | 130-1 | 130-2 |
| --- | --- | --- | --- | --- | --- | --- | --- | --- |
| Reference | Tărlungeanu et al., 2016 |  |  |  |  |  | This study |  |
| SLC7A5 pathogenic variant(s) | c.737C>T<br>p.(Ala246Val) |  |  |  | c.1124C>T<br>p.(Pro375Leu) |  | c.1124C>T,<br>p.(Pro375Leu)<br>Deletion exons 5-10 |  |
| Variant zygosity | Homozygous |  |  |  | Homozygous |  | Compound heterozygous |  |
| Phenotypic characteristics |  |  |  |  |  |  |  |  |
| Gender | F | M | F | M | M | M | F | F |
| Growth retardation | n/a | n/a | n/a | n/a | n/a | n/a | + | + |
| HC at birth (SD) | n/a | -2/-3 | n/a | n/a | -3 | -3 | -3 | -2.5 |
| HC at latest examination (SD) | -5 | -2.3 | -5.5 | -5 | n/a | n/a | -7 | -6 |
| GDD (motor, speech, cognitive) | + | + | + | + | + | + | + | + |
| Seizures (onset) | - | - | - | n/a | +(1y) | +(6mo) | +(3y) | +(6y) |
| Autistic features | + | + | + | n/a | + | + | - | + |
| Brain MRI | n/a | Cortical atrophy | Normal | n/a | Thin CC |  | Pontocerebellar and CC hypoplasia |  |

HC: head circumference, SD: standard deviation, GDD: global developmental delay, n/a: not available, CC: corpus callosum

## REFERENCES

1. Afenjar, A., Moutard, M.-L., Doummar, D., Guët, A., Rabier, D., Vermersch, A.-I., Mignot, C., Burglen, L., Heron, D., Thioulouse, E., et al. (2007). Early neurological phenotype in 4 children with biallelic PRODH mutations. Brain Dev 29, 547–552. https://doi.org/10.1016/j.braindev.2007.01.008.

2. Aon, M.A., Cortassa, S., and O’Rourke, B. (2010). Redox-optimized ROS balance: a unifying hypothesis. Biochim Biophys Acta 1797, 865–877. https://doi.org/10.1016/j.bbabio.2010.02.016.

3. Blanquie, O., Yang, J.-W., Kilb, W., Sharopov, S., Sinning, A., and Luhmann, H.J. (2017). Electrical activity controls area-specific expression of neuronal apoptosis in the mouse developing cerebral cortex. Elife 6, e27696. https://doi.org/10.7554/eLife.27696.

4. Bond, A.M., Ming, G.-L., and Song, H. (2015). Adult Mammalian Neural Stem Cells and Neurogenesis: Five Decades Later. Cell Stem Cell 17, 385–395. https://doi.org/10.1016/j.stem.2015.09.003.

5. Candelario, K.M., Shuttleworth, C.W., and Cunningham, L.A. (2013). Neural stem/progenitor cells display a low requirement for oxidative metabolism independent of hypoxia inducible factor-1alpha expression. J Neurochem 125, 420–429. https://doi.org/10.1111/jnc.12204.

6. Carter, R.J., Morton, J., and Dunnett, S.B. (2001). Motor coordination and balance in rodents. Curr Protoc Neurosci Chapter 8, Unit 8.12. https://doi.org/10.1002/0471142301.ns0812s15.

7. Clark, G.D., Happel, L.T., Zorumski, C.F., and Bazan, N.G. (1992). Enhancement of hippocampal excitatory synaptic transmission by platelet-activating factor. Neuron 9, 1211–1216. https://doi.org/10.1016/0896-6273(92)90078-r.

8. Contreras, X., Amberg, N., Davaatseren, A., Hansen, A.H., Sonntag, J., Andersen, L., Bernthaler, T., Streicher, C., Heger, A., Johnson, R.L., et al. (2021). A genome-wide library of MADM mice for single-cell genetic mosaic analysis. Cell Rep 35, 109274. https://doi.org/10.1016/j.celrep.2021.109274.

9. De Rubeis, S., He, X., Goldberg, A.P., Poultney, C.S., Samocha, K., Cicek, A.E., Kou, Y., Liu, L., Fromer, M., Walker, S., et al. (2014). Synaptic, transcriptional and chromatin genes disrupted in autism. Nature 515, 209–215. https://doi.org/10.1038/nature13772.

10. Dekkers, M.P.J., Nikoletopoulou, V., and Barde, Y.-A. (2013). Cell biology in neuroscience: Death of developing neurons: new insights and implications for connectivity. J Cell Biol 203, 385–393. https://doi.org/10.1083/jcb.201306136.

11. Di Rosa, G., Pustorino, G., Spano, M., Campion, D., Calabrò, M., Aguennouz, M., Caccamo, D., Legallic, S., Sgro, D.L., Bonsignore, M., et al. (2008). Type I hyperprolinemia and proline dehydrogenase (PRODH) mutations in four Italian children with epilepsy and mental retardation. Psychiatr Genet 18, 40–42. https://doi.org/10.1097/YPG.0b013e3282f08a3d.

12. Duran, J., Gruart, A., López-Ramos, J.C., Delgado-García, J.M., and Guinovart, J.J. (2019). Glycogen in Astrocytes and Neurons: Physiological and Pathological Aspects. Adv Neurobiol 23, 311–329. https://doi.org/10.1007/978-3-030-27480-1_10.

13. Dyall, S.C. (2015). Long-chain omega-3 fatty acids and the brain: a review of the independent and shared effects of EPA, DPA and DHA. Front Aging Neurosci 7, 52. https://doi.org/10.3389/fnagi.2015.00052.

14. Elinder, F., and Liin, S.I. (2017). Actions and Mechanisms of Polyunsaturated Fatty Acids on Voltage-Gated Ion Channels. Front Physiol 8, 43. https://doi.org/10.3389/fphys.2017.00043.

15. Fitzgerald, E., Roberts, J., Tennant, D.A., Boardman, J.P., and Drake, A.J. (2021). Metabolic adaptations to hypoxia in the neonatal mouse forebrain can occur independently of the transporters SLC7A5 and SLC3A2. Sci Rep 11, 9092. https://doi.org/10.1038/s41598-021-88757-9.

16. Galler, J.R., Bringas-Vega, M.L., Tang, Q., Rabinowitz, A.G., Musa, K.I., Chai, W.J., Omar, H., Abdul Rahman, M.R., Abd Hamid, A.I., Abdullah, J.M., et al. (2021). Neurodevelopmental effects of childhood malnutrition: A neuroimaging perspective. Neuroimage 231, 117828. https://doi.org/10.1016/j.neuroimage.2021.117828.

17. Gorski, J.A., Talley, T., Qiu, M., Puelles, L., Rubenstein, J.L.R., and Jones, K.R. (2002). Cortical excitatory neurons and glia, but not GABAergic neurons, are produced in the Emx1-expressing lineage. J Neurosci 22, 6309–6314. https://doi.org/20026564.

18. Guyenet, S.J., Furrer, S.A., Damian, V.M., Baughan, T.D., La Spada, A.R., and Garden, G.A. (2010). A simple composite phenotype scoring system for evaluating mouse models of cerebellar ataxia. J Vis Exp 1787. https://doi.org/10.3791/1787.

19. Hammond, J.W., Lu, S.-M., and Gelbard, H.A. (2015). Platelet Activating Factor Enhances Synaptic Vesicle Exocytosis Via PKC, Elevated Intracellular Calcium, and Modulation of Synapsin 1 Dynamics and Phosphorylation. Front Cell Neurosci 9, 505. https://doi.org/10.3389/fncel.2015.00505.

20. Huch, A., Huch, R., Schneider, H., and Rooth, G. (1977). Continuous transcutaneous monitoring of fetal oxygen tension during labour. Br J Obstet Gynaecol 84 *Suppl 1*, 1–39. https://doi.org/10.1111/j.1471-0528.1977.tb16231.x.

21. Iurlaro, R., and Muñoz-Pinedo, C. (2016). Cell death induced by endoplasmic reticulum stress. FEBS J 283, 2640–2652. https://doi.org/10.1111/febs.13598.

22. Kisanuki, Y.Y., Hammer, R.E., Miyazaki, J., Williams, S.C., Richardson, J.A., and Yanagisawa, M. (2001). Tie2-Cre transgenic mice: a new model for endothelial cell-lineage analysis in vivo. Dev Biol 230, 230–242. https://doi.org/10.1006/dbio.2000.0106.

23. Li, J., Shi, M., Ma, Z., Zhao, S., Euskirchen, G., Ziskin, J., Urban, A., Hallmayer, J., and Snyder, M. (2014). Integrated systems analysis reveals a molecular network underlying autism spectrum disorders. Mol Syst Biol 10, 774. https://doi.org/10.15252/msb.20145487.

24. Mason, S. (2017). Lactate Shuttles in Neuroenergetics-Homeostasis, Allostasis and Beyond. Front Neurosci 11, 43. https://doi.org/10.3389/fnins.2017.00043.

25. Moy, S.S., Nadler, J.J., Young, N.B., Perez, A., Holloway, L.P., Barbaro, R.P., Barbaro, J.R., Wilson, L.M., Threadgill, D.W., Lauder, J.M., et al. (2007). Mouse behavioral tasks relevant to autism: phenotypes of 10 inbred strains. Behav Brain Res 176, 4–20. https://doi.org/10.1016/j.bbr.2006.07.030.

26. Murphy, M.P. (2009). How mitochondria produce reactive oxygen species. Biochem J 417, 1–13. https://doi.org/10.1042/BJ20081386.

27. Napolitano, L., Scalise, M., Galluccio, M., Pochini, L., Albanese, L.M., and Indiveri, C. (2015). LAT1 is the transport competent unit of the LAT1/CD98 heterodimeric amino acid transporter. Int J Biochem Cell Biol 67, 25–33. https://doi.org/10.1016/j.biocel.2015.08.004.

28. Nikolić, M., Gardner, H. a. R., and Tucker, K.L. (2013). Postnatal neuronal apoptosis in the cerebral cortex: physiological and pathophysiological mechanisms. Neuroscience 254, 369–378. https://doi.org/10.1016/j.neuroscience.2013.09.035.

29. Nwadike, C., Williamson, L.E., Gallagher, L.E., Guan, J.-L., and Chan, E.Y.W. (2018). AMPK Inhibits ULK1-Dependent Autophagosome Formation and Lysosomal Acidification via Distinct Mechanisms. Molecular and Cellular Biology 38, e00023–18. https://doi.org/10.1128/MCB.00023-18.

30. Onishi, Y., Hiraiwa, M., Kamada, H., Iezaki, T., Yamada, T., Kaneda, K., and Hinoi, E. (2019). Hypoxia affects Slc7a5 expression through HIF-2α in differentiated neuronal cells. FEBS Open Bio 9, 241–247. https://doi.org/10.1002/2211-5463.12559.

31. Parenti, I., Rabaneda, L.G., Schoen, H., and Novarino, G. (2020). Neurodevelopmental Disorders: From Genetics to Functional Pathways. Trends in Neurosciences 43, 608–621. https://doi.org/10.1016/j.tins.2020.05.004.

32. Philips, T., and Rothstein, J.D. (2017). Oligodendroglia: metabolic supporters of neurons. J Clin Invest 127, 3271–3280. https://doi.org/10.1172/JCI90610.

33. Richards, S., Aziz, N., Bale, S., Bick, D., Das, S., Gastier-Foster, J., Grody, W.W., Hegde, M., Lyon, E., Spector, E., et al. (2015). Standards and guidelines for the interpretation of sequence variants: a joint consensus recommendation of the American College of Medical Genetics and Genomics and the Association for Molecular Pathology. Genet Med 17, 405–423. https://doi.org/10.1038/gim.2015.30.

34. Riggs, A.C., Bernal-Mizrachi, E., Ohsugi, M., Wasson, J., Fatrai, S., Welling, C., Murray, J., Schmidt, R.E., Herrera, P.L., and Permutt, M.A. (2005). Mice conditionally lacking the Wolfram gene in pancreatic islet beta cells exhibit diabetes as a result of enhanced endoplasmic reticulum stress and apoptosis. Diabetologia 48, 2313–2321. https://doi.org/10.1007/s00125-005-1947-4.

35. Robb, E.L., Hall, A.R., Prime, T.A., Eaton, S., Szibor, M., Viscomi, C., James, A.M., and Murphy, M.P. (2018). Control of mitochondrial superoxide production by reverse electron transport at complex I. J Biol Chem 293, 9869–9879. https://doi.org/10.1074/jbc.RA118.003647.

36. Rock, K.D., and Patisaul, H.B. (2018). Environmental Mechanisms of Neurodevelopmental Toxicity. Curr Environ Health Rep 5, 145–157. https://doi.org/10.1007/s40572-018-0185-0.

37. Ross, E.J., Graham, D.L., Money, K.M., and Stanwood, G.D. (2015). Developmental Consequences of Fetal Exposure to Drugs: What We Know and What We Still Must Learn. Neuropsychopharmacology 40, 61–87. https://doi.org/10.1038/npp.2014.147.

38. Sinclair, L.V., Rolf, J., Emslie, E., Shi, Y.-B., Taylor, P.M., and Cantrell, D.A. (2013). Control of amino-acid transport by antigen receptors coordinates the metabolic reprogramming essential for T cell differentiation. Nat Immunol 14, 500–508. https://doi.org/10.1038/ni.2556.

39. Southwell, D.G., Paredes, M.F., Galvao, R.P., Jones, D.L., Froemke, R.C., Sebe, J.Y., Alfaro-Cervello, C., Tang, Y., Garcia-Verdugo, J.M., Rubenstein, J.L., et al. (2012). Intrinsically determined cell death of developing cortical interneurons. Nature 491, 109–113. https://doi.org/10.1038/nature11523.

40. Stankovic, I.N., and Colak, D. (2022). Prenatal Drugs and Their Effects on the Developing Brain: Insights From Three-Dimensional Human Organoids. Frontiers in Neuroscience 16.

41. Takahara, T., Amemiya, Y., Sugiyama, R., Maki, M., and Shibata, H. (2020). Amino acid-dependent control of mTORC1 signaling: a variety of regulatory modes. Journal of Biomedical Science 27, 87. https://doi.org/10.1186/s12929-020-00679-2.

42. Tărlungeanu, D.C., Deliu, E., Dotter, C.P., Kara, M., Janiesch, P.C., Scalise, M., Galluccio, M., Tesulov, M., Morelli, E., Sonmez, F.M., et al. (2016). Impaired Amino Acid Transport at the Blood Brain Barrier Is a Cause of Autism Spectrum Disorder. Cell 167, 1481–1494.e18. https://doi.org/10.1016/j.cell.2016.11.013.

43. Vilchez, D., Ros, S., Cifuentes, D., Pujadas, L., Vallès, J., García-Fojeda, B., Criado-García, O., Fernández-Sánchez, E., Medraño-Fernández, I., Domínguez, J., et al. (2007). Mechanism suppressing glycogen synthesis in neurons and its demise in progressive myoclonus epilepsy. Nat Neurosci 10, 1407–1413. https://doi.org/10.1038/nn1998.

44. Wong, F.K., Bercsenyi, K., Sreenivasan, V., Portalés, A., Fernández-Otero, M., and Marín, O. (2018). Pyramidal cell regulation of interneuron survival sculpts cortical networks. Nature 557, 668–673. https://doi.org/10.1038/s41586-018-0139-6.

45. Wortel, I.M.N., van der Meer, L.T., Kilberg, M.S., and van Leeuwen, F.N. (2017). Surviving Stress: Modulation of ATF4-Mediated Stress Responses in Normal and Malignant Cells. Trends Endocrinol Metab 28, 794–806. https://doi.org/10.1016/j.tem.2017.07.003.

46. Xia, K., Guo, H., Hu, Z., Xun, G., Zuo, L., Peng, Y., Wang, K., He, Y., Xiong, Z., Sun, L., et al. (2014). Common genetic variants on 1p13.2 associate with risk of autism. Mol Psychiatry 19, 1212–1219. https://doi.org/10.1038/mp.2013.146.

47. Ye, Z., Wang, S., Zhang, C., and Zhao, Y. (2020). Coordinated Modulation of Energy Metabolism and Inflammation by Branched-Chain Amino Acids and Fatty Acids. Frontiers in Endocrinology 11.

48. Yu, H., Villanueva, N., Bittar, T., Arsenault, E., Labonté, B., and Huan, T. (2020). Parallel metabolomics and lipidomics enables the comprehensive study of mouse brain regional metabolite and lipid patterns. Analytica Chimica Acta 1136, 168–177. https://doi.org/10.1016/j.aca.2020.09.051.

49. Zhang, L., Ou, J., Xu, X., Peng, Y., Guo, H., Pan, Y., Chen, J., Wang, T., Peng, H., Liu, Q., et al. (2015). AMPD1 functional variants associated with autism in Han Chinese population. Eur Arch Psychiatry Clin Neurosci 265, 511–517. https://doi.org/10.1007/s00406-014-0524-6.

50. Zhang, S., Lin, X., Hou, Q., Hu, Z., Wang, Y., and Wang, Z. (2021). Regulation of mTORC1 by amino acids in mammalian cells: A general picture of recent advances. Animal Nutrition 7, 1009–1023. https://doi.org/10.1016/j.aninu.2021.05.003.

51. Zong, H., Espinosa, J.S., Su, H.H., Muzumdar, M.D., and Luo, L. (2005). Mosaic analysis with double markers in mice. Cell 121, 479–492. https://doi.org/10.1016/j.cell.2005.02.012.

